# Immune Cell Profiling Reveals MAIT and Effector Memory CD4+ T Cell Recovery Link to Control of Cytomegalovirus Reactivation after Stem Cell Transplant

**DOI:** 10.1101/2021.08.09.455593

**Authors:** Lauren Stern, Helen M. McGuire, Selmir Avdic, Barbara Fazekas de St Groth, David Gottlieb, Allison Abendroth, Emily Blyth, Barry Slobedman

## Abstract

Human cytomegalovirus (HCMV) reactivation is a major opportunistic infection after allogeneic haematopoietic stem cell transplantation and has a complex relationship with post-transplant immune reconstitution. Here, we used mass cytometry to comprehensively define global patterns of innate and adaptive immune cell reconstitution at key phases of HCMV reactivation (before detection, initial detection, peak and near resolution) in the first 100 days post-transplant. In addition to identifying patterns of immune reconstitution in those with or without HCMV reactivation, we found mucosal-associated invariant T (MAIT) cell levels at the initial detection of HCMV DNAemia distinguished patients who subsequently developed low-level versus high-level HCMV reactivation. In addition, early recovery of effector-memory CD4^+^ T cells distinguished low-level and high-level reactivation. Our data describe distinct immune signatures that emerged with HCMV reactivation post-HSCT, and highlight MAIT cell levels at the initial detection of reactivation as a potential prognostic marker to guide clinical decisions regarding pre-emptive therapy.

## INTRODUCTION

Human cytomegalovirus (HCMV) reactivation is a major complication after allogeneic haematopoietic stem cell transplantation (HSCT) (Ljungman et al., 2011; Stern et al., 2019). It is associated with significant morbidity, increased non-relapse mortality, microbial superinfections and acute graft-versus-host disease (aGvHD) (Cantoni et al., 2010; Giménez et al., 2019; Green et al., 2016; Teira et al., 2016). A dose-response relationship between the magnitude of HCMV viraemia and risk of mortality has been reported (Green et al., 2016). While antiviral prophylaxis or pre-emptive therapy (initiated following detection of HCMV DNAemia) can reduce the incidence of end-organ HCMV disease, commonly used antiviral drugs such as ganciclovir and foscarnet carry significant toxicities (Zavras et al., 2020). Improved risk stratification methods are needed to enable early intervention in patients who will develop high-level HCMV viraemia, while minimising unnecessary exposure to antivirals in those who experience low-level, self-limiting HCMV reactivation.

Immune reconstitution after HSCT has a close relationship with HCMV reactivation (Blyth et al., 2016). Measurements of immune reconstitution, such as HCMV-specific T cell responses, may be integrated alongside viral load monitoring to guide the targeting of antiviral therapy (Avetisyan et al., 2007; Camargo et al., 2019; Duke et al., 2020; El Haddad et al., 2019; Gabanti et al., 2021; Green et al., 2012; Lilleri et al., 2012; Navarro et al., 2016; Tey et al., 2013). High-dimensional technologies such as mass cytometry (CyTOF) (Bandura et al., 2009; Ornatsky et al., 2010), which utilises antibodies conjugated to heavy-metal isotopes to allow simultaneous assessment of >40 single-cell markers, have the potential to aid in identification of clinically important immune signatures and improve our understanding of the complex relationship between HCMV reactivation, immune reconstitution and clinical outcome (Stern et al., 2018).

Herein, we used CyTOF to investigate the dynamics of innate and adaptive immune cell reconstitution in the first 100 days post-transplant in a cohort of 35 adult allogeneic HSCT patients who experienced high-level, low-level or no HCMV reactivation. Comprehensive immunophenotyping was performed on peripheral blood samples from four time-points aligning with the kinetics of HCMV DNAemia (prior to detection, initial detection, peak, and near the resolution). Our results identified MAIT cells and effector-memory (EM) CD4^+^ T cells as key features distinguishing patients with low-level and high-level HCMV reactivation, and provide a detailed analysis of patterns of immune recovery accompanying different phases of HCMV reactivation in the first 100 days post-HSCT.

## RESULTS

### Patient characteristics

We studied 35 allogeneic HSCT recipients in the first 100 days post-transplant (**Figure 1A**). Patients characteristics are outlined in **Table 1**. Of the 24 patients at risk of HCMV reactivation (i.e. HCMV-seropositive donor (D+) and/or recipient (R+)), HCMV reactivation was detected in 19 (79.2%) patients, at a median 29 (12-46) days post-HSCT. Two patterns of HCMV reactivation were observed (**Figure 1A**); low-level reactivation (LR; <250 peak HCMV DNA copies/mL; n=6) and high-level reactivation (HR; >830 peak copies/mL; n=13). We thus retrospectively divided the cohort into four study groups (**Figure 1A**): LR (low-level HCMV reactivation; n=6), HR (high-level HCMV reactivation; n=13), SP-NR (HCMV-seropositive D+ and/or recipient R+, with no documented HCMV reactivation; n=5), and SN (D-/R-HCMV-seronegative, with no HCMV infection; n=11). Peak HCMV DNAemia levels were significantly higher in HR (median 3873 (836-52740) copies/mL) compared to LR (150 (150-244) copies/mL; p<0.0001) patients (**Figure S1A**). No patient was diagnosed with HCMV end-organ disease in the follow-up period.

**Figure 1.**
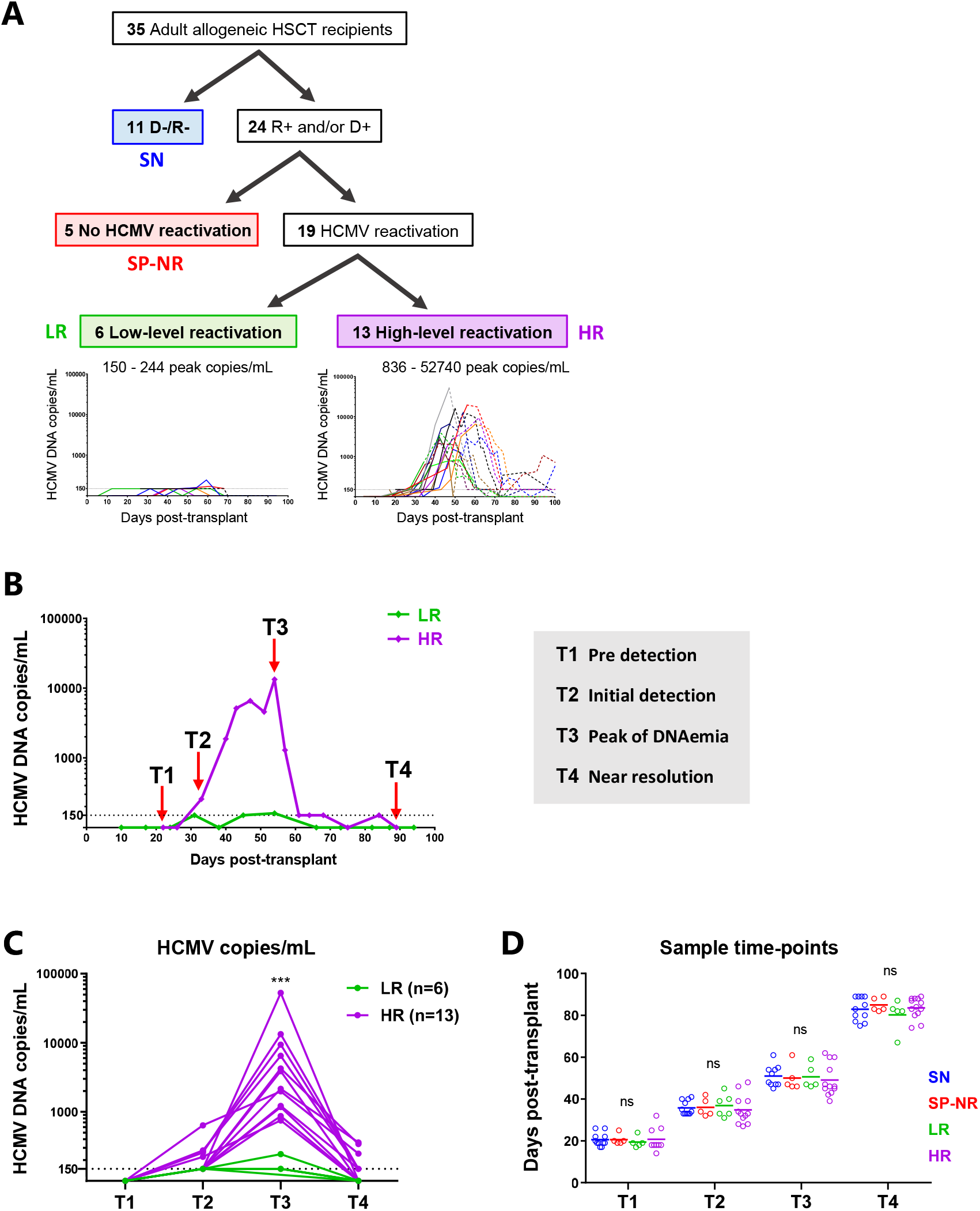
Study design. (**A**) HSCT recipients were divided into four groups: SN (HCMV-seronegative D-/R-; blue), SP-NR (HCMV-seropositive R+ and/or D+, with no detected HCMV DNAemia in the first 100 days post-HSCT; red), LR (patients who developed low-level (<250 peak copies/mL) HCMV DNAemia; green), and HR (patients who developed high-level (>830 peak copies/mL) HCMV DNAemia; purple). Graphs show HCMV DNA copies/mL plasma in the first 100 days post-transplant for LR (left) and HR (right). Each line represents one patient. The point of antiviral therapy initiation for each patient is indicated by the shift from solid line (pre-therapy) to dotted line (after therapy initiation). The horizontal dotted line at 150 copies/mL indicates the lower limit of quantitation (LLQ). (**B**) Representative HCMV reactivation profiles from one HR and one LR patient depicting the four time-points analysed with mass cytometry. Peripheral blood mononuclear cells (PBMC) collected prior to detection (T1), at the initial detection (T2), at the peak (T3) and near the resolution (T4) of HCMV DNAemia post-HSCT were analysed. The horizontal dotted line indicates the LLQ (150 HCMV DNA copies/mL). Samples from corresponding days post-transplant were analysed from HSCT recipients without HCMV reactivation. (**C**) HCMV DNA plasma copy load at each study time-point from patients with HCMV reactivation. Each line represents a patient. The horizontal dotted line at 150 copies/mL indicates the LLQ. Two-tailed Mann-Whitney test comparing HR and LR per time point (*** p<0.001). (**D**) Days post-transplant for all PBMC samples analysed with mass cytometry. Lines indicate the mean. One-way ANOVA per time-point with Fisher’s Least Significant Difference test (ns, not significant). SN, HCMV-seronegative; SP-NR, seropositive no reactivation; LR, low-level HCMV reactivation; HR, high-level HCMV reactivation. D, donor; R, recipient; HCMV (human cytomegalovirus); HSCT (haematopoietic stem cell transplant).

**Table 1.**
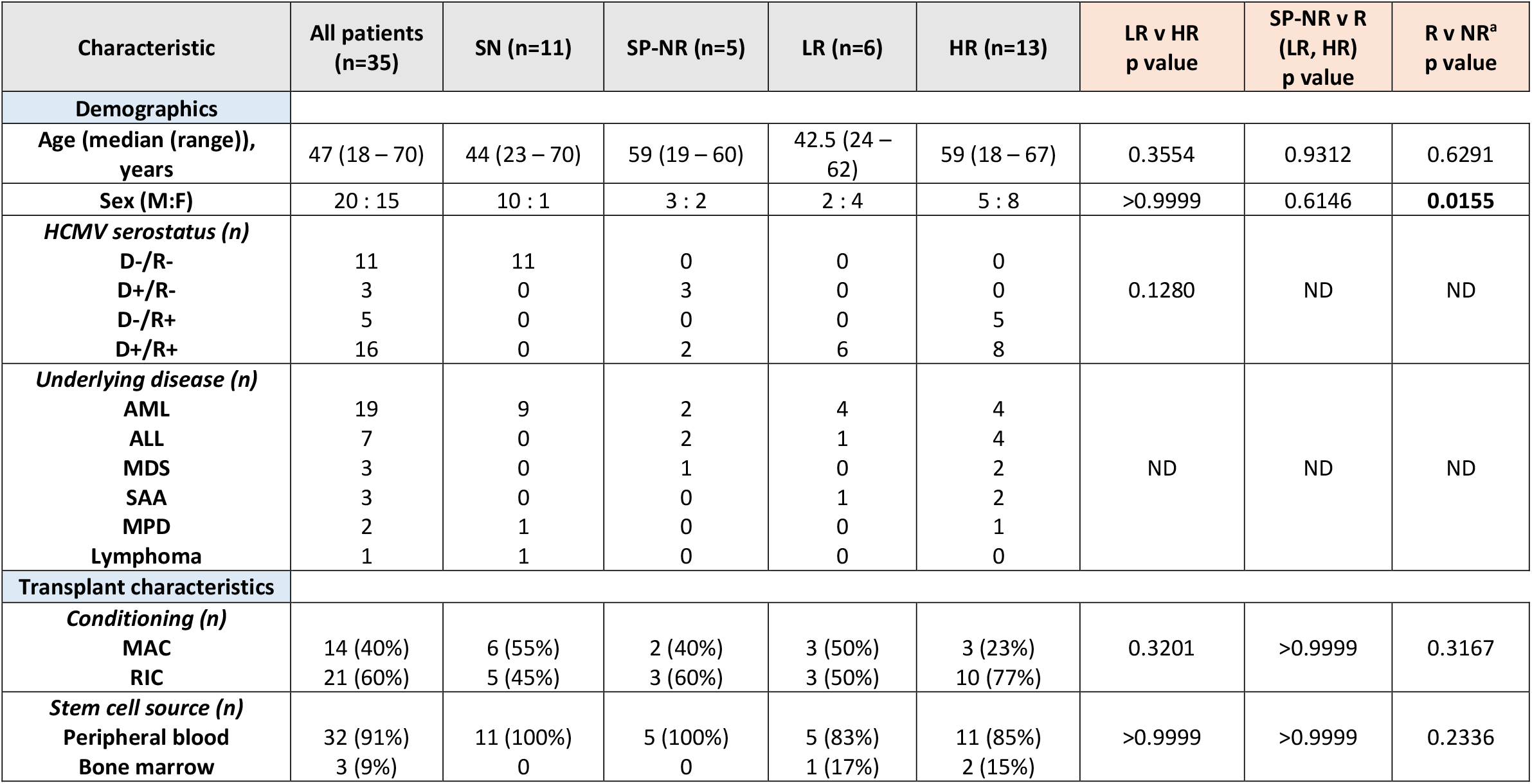
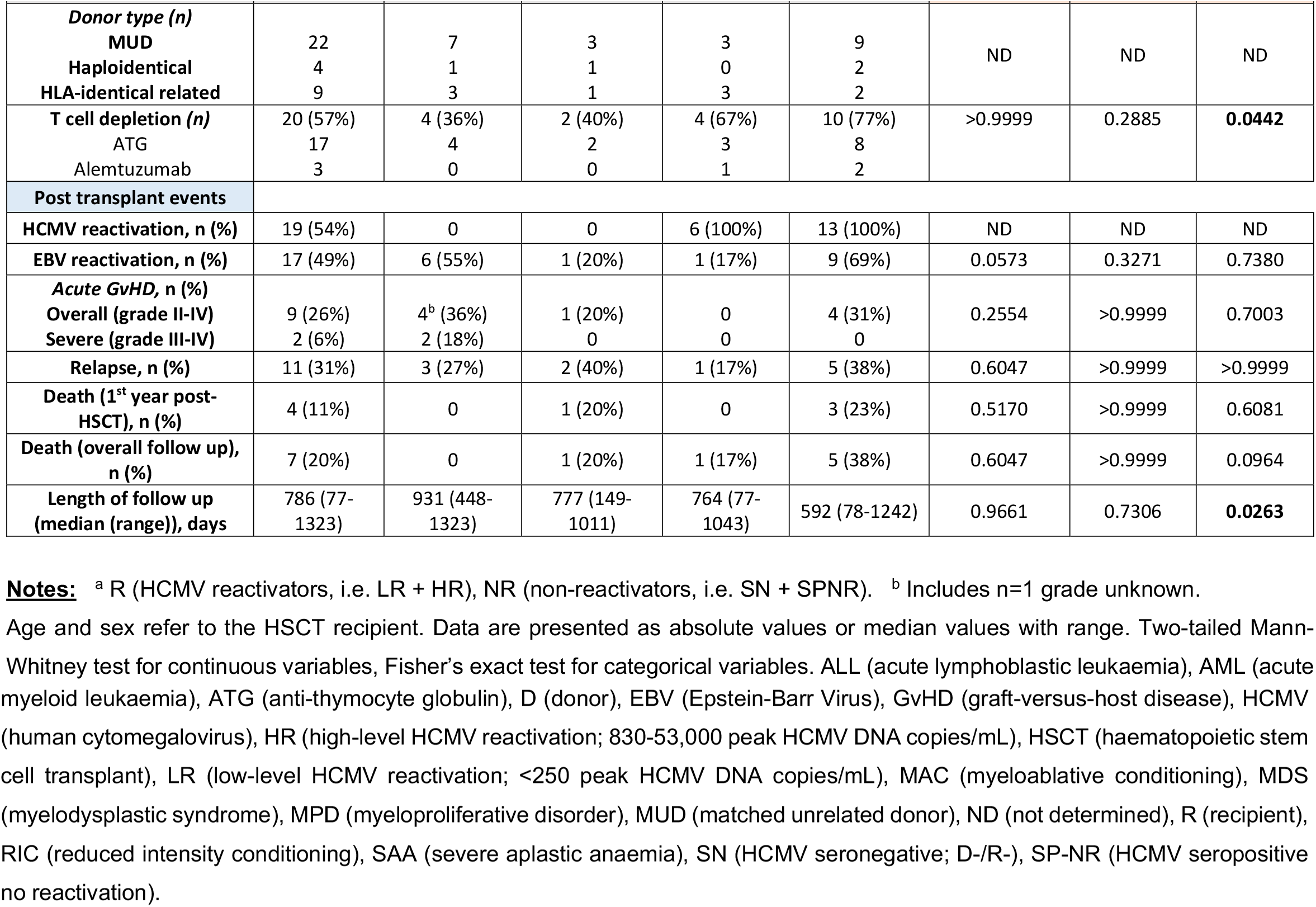
Characteristics of study subjects.

Overall survival was 88.6% (31/35) in first year post-transplant (**Table 1**). Both patients who died in the first 100 days, and 3/4 patients who died in the first year post-transplant, were HR patients (additional patient was SP-NR). There was a higher incidence of EBV reactivation in HR vs. LR groups (9/13 (69.2%) vs. 1/6 (16.7%); p=0.0573). Incidence of aGvHD grades II-IV was also higher in HR (4/13) than LR (0/6) (**Table 1**).

### Features of HCMV reactivation

The median day and magnitude of first detected HCMV DNAemia did not differ significantly between LR and HR (**Table S1**). However, HR had a longer duration of HCMV DNAemia than LR (median 60 (28-121) vs. 21.5 (7-49) days, respectively; p=0.0180), and the log_10_ area under the curve of HCMV DNA copies/mL (a measure of HCMV antigen exposure) in the first 100 days post-HSCT (AUC_0-100_) was significantly higher in HR than LR (**Figure S1B**). All (13/13) HR patients required antiviral therapy to control HCMV reactivation, whereas a majority of LR patients (5/6) had self-limiting HCMV DNAemia (p=0.0005). The one LR patient who received antiviral therapy had low HCMV DNAemia levels consistent with the other LR patients (**Figure 1A**). Resolution of HCMV reactivation by day 100 post-HSCT was achieved in 6/6 (100%) LR patients, compared to 6/13 (46.2%) HR patients (p=0.0436).

### Increased CD8^+^ T cell number and inverted CD4:CD8 ratio after HCMV reactivation

To comprehensively describe immune reconstitution at key time-points during HCMV reactivation post-HSCT, we employed CyTOF to analyse PBMC samples collected prior to detection (T1), at initial detection (T2), at peak (T3) and near resolution (T4) of HCMV DNAemia in the first 100 days post-HSCT (**Figure 1B; Table S2**). HCMV DNA copy numbers per time-point are shown in **Figure 1C**. Samples from matched days post-transplant were analysed in parallel from HSCT recipients who did not experience HCMV reactivation (SN and SP-NR) (**Figure 1D**). PBMCs were stained with a panel of 36 heavy metal isotope-tagged antibodies (**Table S3**) allowing identification of 77 innate and adaptive immune subsets at each time-point (see **Table S4; Table S5; Figure S2**).

There were no significant differences in the absolute counts of major PBMC subsets between the groups at T1 (**Figure 2A**). Subsequently, faster numerical recovery of CD8^+^ T cells, CD3^+^CD56^+^ cells and γδ T cells was observed in patients with HCMV reactivation. Recovery dynamics of innate populations including monocytes and NK cells were similar in all patient groups (**Figure 2A**). LR patients demonstrated the fastest reconstitution of CD8^+^ T cells, CD3^+^CD56^+^ cells and γδ T cells, culminating in significantly higher γδ T cell and CD3^+^CD56^+^ cell counts at T4. Significant increases in NK cells, CD3^+^CD56^+^ cells, γδ T cells, CD8^+^ T cells and CD4^+^ T cells between T1 and T2 were evident only in LR patients. HR patients displayed the slowest CD4^+^ T cell recovery, but showed clear increases in B cells and CD8^+^ T cells at T3. The CD4:CD8 T cell ratio became inverted in HCMV reactivators, diverging at T2 in LR and T3 in HR, while patients without HCMV reactivation maintained a CD4 bias across the time-points (**Figure 2B**).

**Figure 2.**
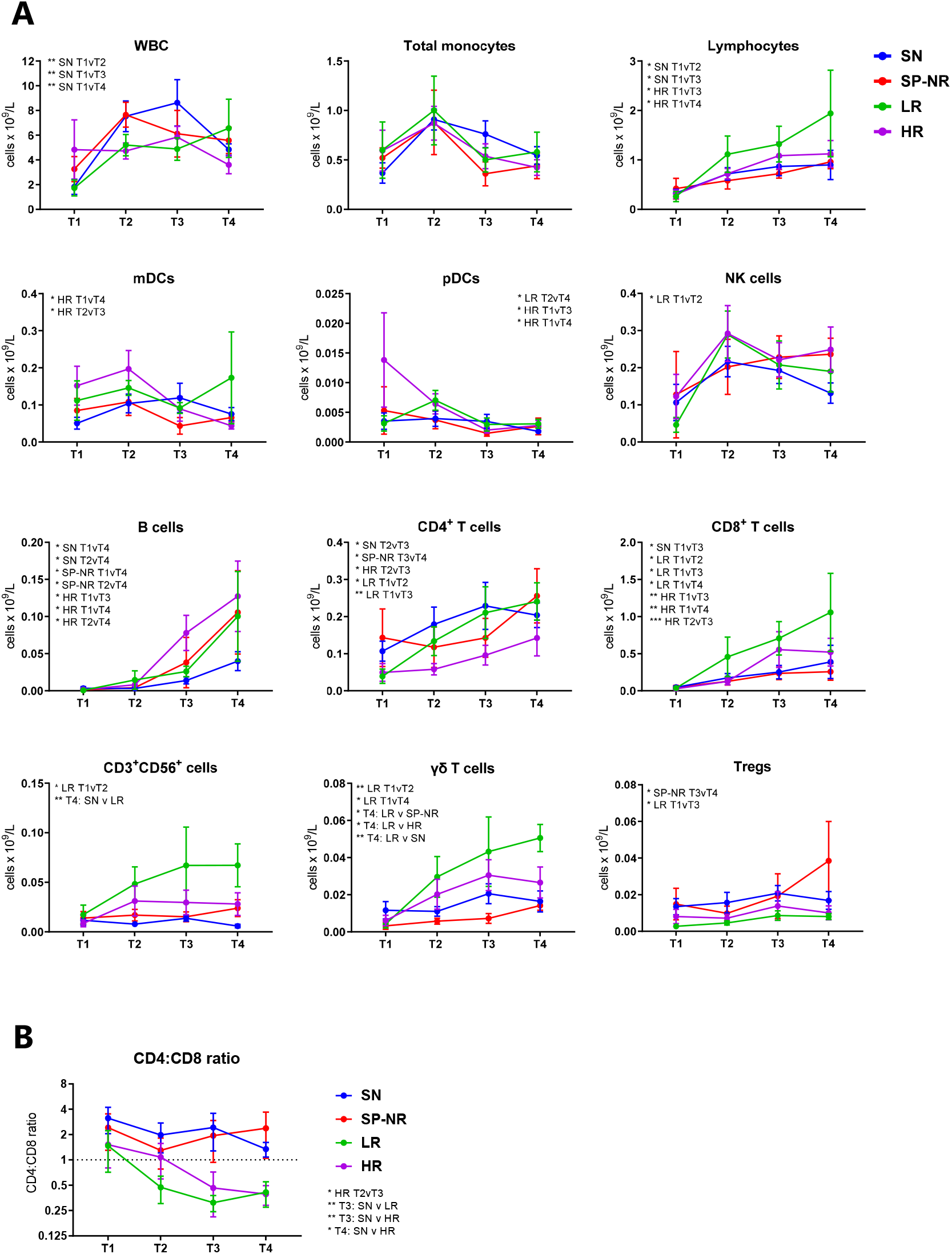
Reconstitution kinetics of peripheral blood mononuclear cell populations in the first 100 days post-HSCT. (**A**) Absolute blood counts (×10^9^/L) of major immune cell subsets over the four study time-points in HSCT recipients with (LR, HR) or without (SN, SP-NR) HCMV reactivation. T1 is prior to the detection of HCMV reactivation; T2, at the initial detection of HCMV reactivation; T3, the peak; T4, near the resolution of HCMV reactivation. Blood samples from corresponding days post-HSCT were assessed in parallel from HSCT recipients without HCMV reactivation. Statistically significant comparisons are listed on the figure for each cell type (mixed-effects model with Tukey’s multiple comparisons test; * p<0.05, ** p<0.01, *** p<0.001). ‘v’ indicates time-points or groups being compared. (**B**) CD4:CD8 T cell ratio. Mixed-effects model with Tukey’s multiple comparisons test (* p<0.05, ** p<0.01, *** p<0.001). Significant comparisons are listed in text on the figure. All graphs show mean ± SEM. SN, HCMV-seronegative (n=11); SP-NR, seropositive no reactivation (n=5); LR, low-level HCMV reactivation (n=6); HR, high-level HCMV reactivation (n=13). mDC, myeloid dendritic cell; NK, natural killer; pDC, plasmacytoid dendritic cell; Treg, T regulatory cell; WBC, white blood cells.

### Distinct pattern of immune reconstitution in patients with HCMV reactivation

To detail the immune profile at each time-point, we used significance analysis of microarrays (SAM) to compare all immune subsets analysed between patients with or without HCMV reactivation, examining both cell proportions and absolute counts. Consistent with **Figure 2**, no significant differences in cell proportions (**Figure 3A**) or absolute counts (**Figure 3B**) were identified between ‘reactivators’ (LR and HR) and ‘non-reactivators’ (SN and SP-NR) at T1. However, multiple cell phenotypes indicative of immune activation were enriched in reactivators at T2 (**Figure 3A**), including elevated percentages of CD86^+^ monocyte subsets and myeloid dendritic cells (mDC), HLA-DR^+^CD38^+^ total and EM CD4^+^ and CD8^+^ T cells, and higher EM, PD-1^+^, and ICOS^+^ percentages in the CD4^+^ T cell compartment. In contrast, few differences in absolute cell counts were observed between reactivators and non-reactivators at T2, although central-memory (CM) and CD27^+^CD4^+^ T cell numbers appeared lower in reactivators (**Figure 3B**).

**Figure 3.**
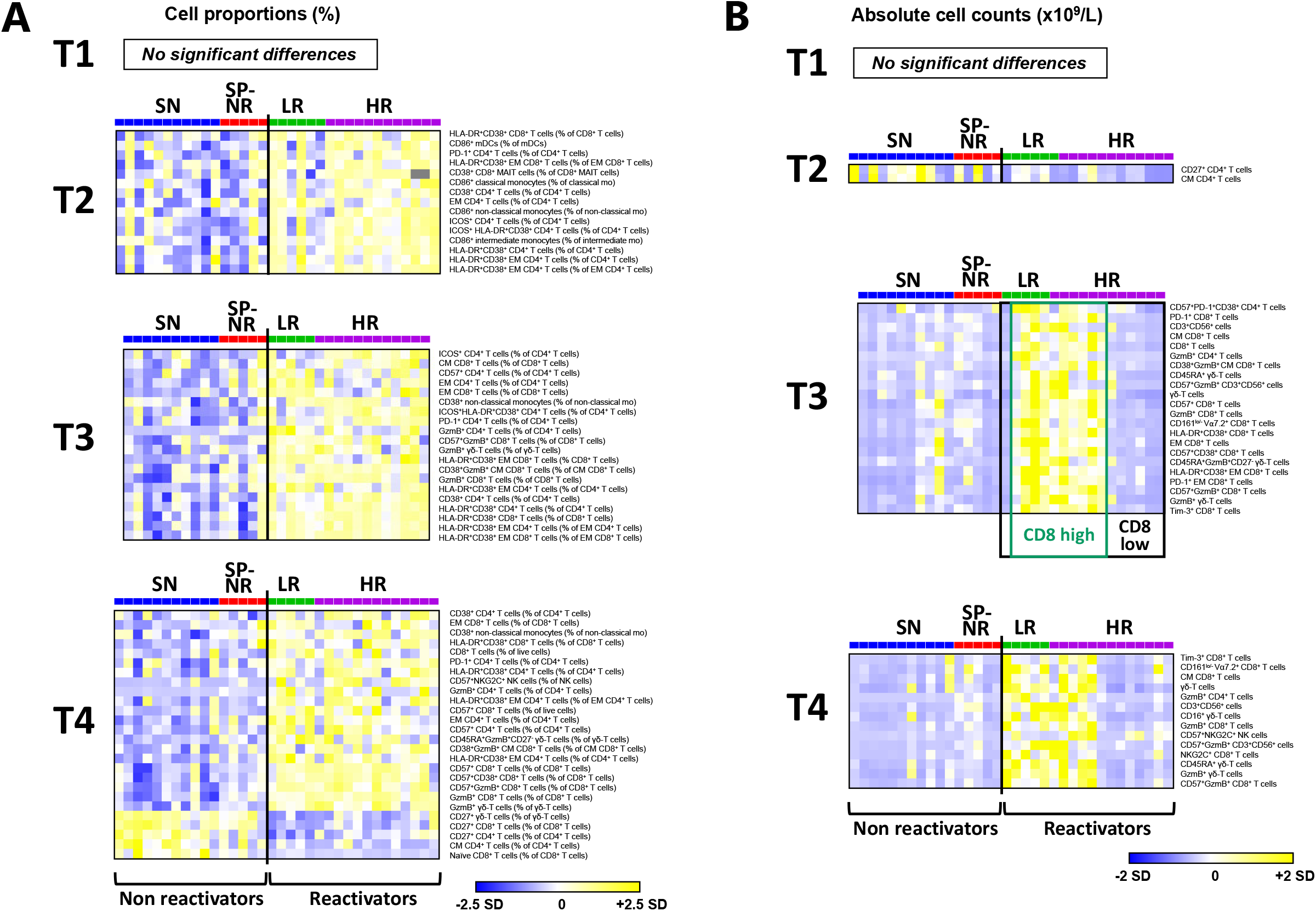
Distinct immune reconstitution pattern in HSCT recipients with HCMV reactivation. (**A**) Heat map rows display percentages of immune cell subsets that were significantly different between HSCT recipients who did (LR, HR) or did not (SN, SP-NR) experience HCMV reactivation in the first 100 days post-transplant. Percentages are expressed as percent of live cells and/or percent of parent subset. Significance was determined using two-class unpaired significance analysis of microarrays (SAM) involving 88 cell subsets per time-point (see **Table S4**). Heat maps are coloured by the Z-score normalised per row (cell subset). Grey colour indicates missing data (too few cells in parent gate to accurately gate subpopulation). Each column represents a patient. (**B**) Heat map rows display cell subsets (absolute cell counts; x10^9^/L blood) that differ significantly between HCMV reactivators (LR, HR) and non-reactivators (SN, SP-NR) as determined by two-class unpaired SAM analysis involving 77 populations per time-point (see **Table S5**). Heat maps are coloured by the Z-score normalised per row (cell subset). Each column represents a patient. Patients with HCMV reactivation were subsequently stratified into ‘CD8^high^’ (dark green box) or ‘CD8^low^’ (black box) groups according to the presence or absence of the elevated CD8^+^ T cell dominated profile observed at T3. See **Figure S3** for a schematic representation of cell subsets shown in **Figure 3A-B**. T1 is prior to the detection of HCMV reactivation; T2, at the initial detection of HCMV reactivation; T3, the peak; T4, near the resolution of HCMV reactivation. Naïve (CD45RA^+^ CD45RO^-^ CCR7^+^ CD27^+^), CM (central memory; CD45RA^-^ CD45RO^+^ CCR7^+^ CD27^+^), EM (effector memory; CD45RA^-^ CD45RO^+^ CCR7^-^CD27^-^), Mo (monocyte). SN, HCMV-seronegative (blue; n=11); SP-NR, seropositive no reactivation (red; n=5); LR, low-level HCMV reactivation (green; n=6); HR, high-level HCMV reactivation (purple; n=13); SD, standard deviation.

At T3 and T4, reactivators maintained a distinct immune profile from non-reactivators, with many shared features evident between LR and HR patients (**Figure 3A**). This included memory T cell populations at significantly higher percentages in reactivators. Elevated percentages of CD4^+^, CD8^+^ and γδ T cells expressing granzyme-B (GzmB) and/or CD57 emerged at T3 in reactivators and remained elevated at T4; these markers indicative of cytotoxic potential have previously been noted to be expressed by HCMV-specific T cells (Raeiszadeh et al., 2015; Scheinberg et al., 2009; Suessmuth et al., 2015; van Leeuwen et al., 2004; Yeh et al., 2021). A majority (15/19; 79%) of subsets detected at significantly higher percentages in reactivators at T3 remained so at T4 (**Figure 3A; Figure S3A**). At T4, significantly lower percentages of CD27-expressing T cells, including naïve CD8^+^ T cells and CM CD4^+^ T cells, as well as higher proportions of CD57^+^NKG2C^+^ NK cells and CD57^+^CD8^+^ T cell subsets, were evident in reactivators (**Figure 3A**). These results highlight multiple phenotypic differences in immune reconstitution over the course of HCMV reactivation, culminating in a markedly distinct T cell dominated immune profile at T4.

### Elevated CD8^+^ T cell dominated signature at the peak of HCMV reactivation in a subset of reactivators

Comparison of absolute cell counts between reactivators and non-reactivators exposed two patterns of quantitative immune recovery within the reactivator group at T3 and T4 (**Figure 3B**). A CD8^+^ T cell dominated signature (‘CD8^high^’) comprising significantly higher absolute numbers of activated and memory CD8^+^ T cells, as well as γδ T cell, CD3^+^CD56^+^ cell and CD4^+^ T cell subsets, was evident at T3 in most (4/5) of LR patients and half (6/12) of HR patients (**Figure 3B**). At T4, this elevated quantitative immune profile contained 11/22 populations seen at T3, as well as CD57^+^NKG2C^+^ NK cells, NKG2C^+^CD8^+^ T cells and CD16^+^ γδ T cells (**Figure 3B; Figure S3B-C**).

Direct comparison of all cell-subset counts between reactivators with the ‘CD8^high^’ or ‘CD8^low^’ signature at the peak of HCMV reactivation revealed numerous CD8^+^ T cell and γδ T cell subsets that were significantly higher in CD8^high^ reactivators at T3 (**Figure S4A**). CD8^+^ T cell frequencies did not differ between CD8^high^ and CD8^low^ patients before the detection of HCMV reactivation (T1), but CD8^high^ patients displayed faster recovery of total CD8^+^ T cell absolute counts and percentages over the course of HCMV reactivation (**Figure 4A-B**). Total CD8^+^ T cell counts at T3 were significantly higher in CD8^high^ reactivators compared to CD8^low^ reactivators (median 0.7634 vs. 0.07837 ×10^9^/L, respectively; p=0.0004) and non-reactivators (median 0.1291 ×10^9^/L; p=0.0027), but there was no significant difference in CD8^+^ T cell count between non-reactivators and CD8^low^ reactivators (**Figure 4A; Figure S4B**). These findings highlight differences in the response to HCMV reactivation and indicate that enhanced CD8^+^ T cell recovery is not a universal feature in HSCT recipients at the peak of HCMV reactivation.

**Figure 4.**
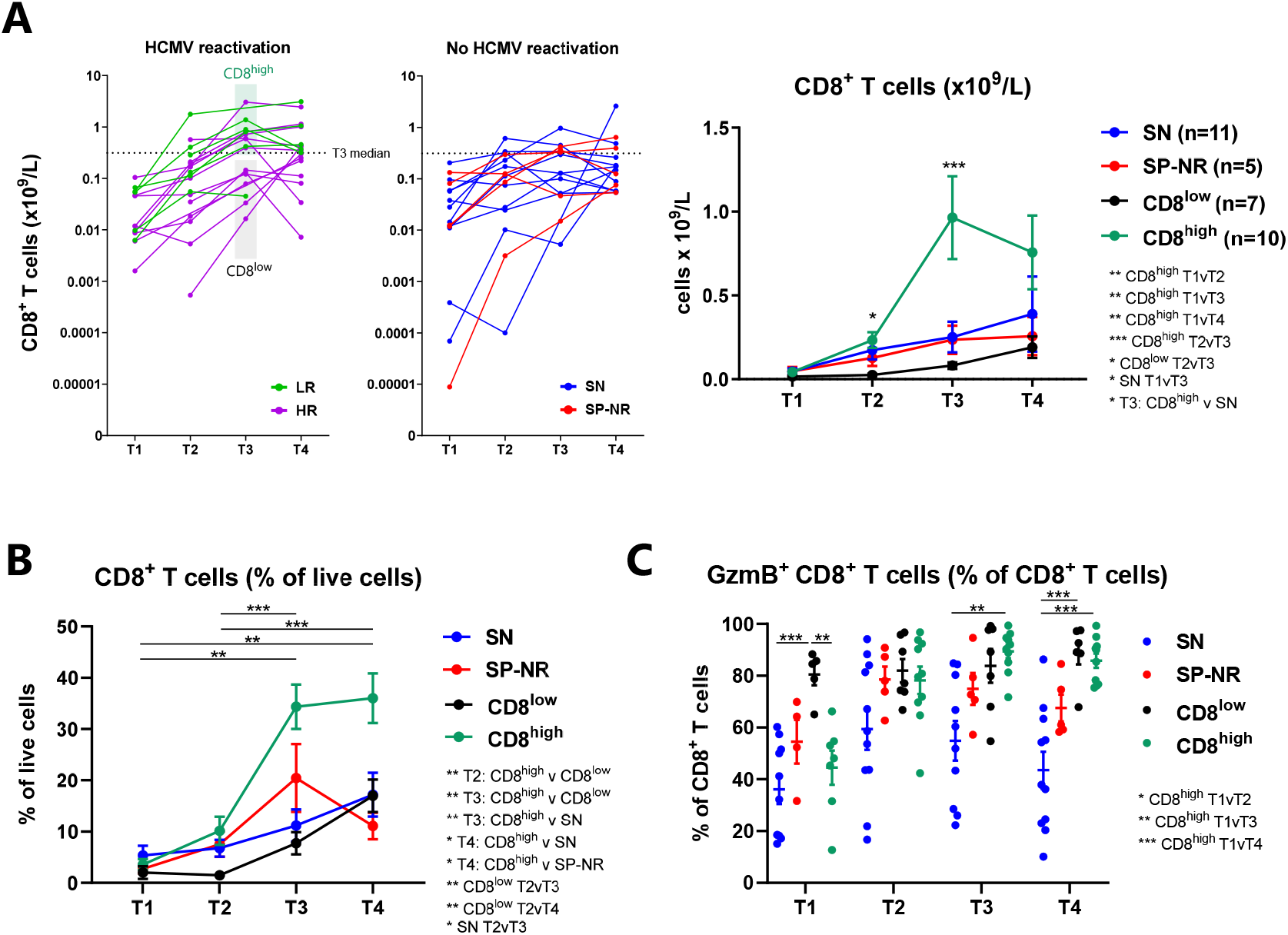
Emergence of a CD8^+^ T cell dominated immune profile in a subset of HSCT recipients with HCMV reactivation. (**A**) Absolute CD8^+^ T cell counts (x10^9^/L) in HSCT recipients with or without HCMV reactivation (as in Figure 2A). Each line represents one patient. The horizontal black dotted line shows the median CD8^+^ T cell count across all patients at T3 (0.3116 ×10^9^/L). Note that the Y-axis displays a log_10_ scale. HSCT recipients with HCMV reactivation were partitioned in ‘CD8^high^’ (n=10) and ‘CD8^low^’ (n=7) groups according to the presence or absence of a CD8^+^ T cell dominated immune signature identified at the peak of HCMV reactivation (T3) in **Figure 3B**. Graph on right shows mean ± SEM for each group; the two HCMV reactivators (n=1 LR, n=1 HR) without a PBMC sample available at T3 are not included. Statistical significance was evaluated using mixed-effects models with Tukey’s multiple comparisons test (* p<0.05, ** p<0.01, *** p<0.001). Asterisks on figure at T2 and T3 indicate significant differences between ‘CD8^high^’ (dark green) and ‘CD8^low^’ (black) groups. Additional significant comparisons are listed in text adjacent to the graph (‘v’ indicates time-points or groups being compared). (**B**) CD8^+^ T cells (as a percentage of total live cells) with mean ± SEM in CD8^high^ reactivators (dark green, n=10), CD8^low^ reactivators (black, n=7) and patients without HCMV reactivation (SN (blue, n=11), SP-NR (red, n=5)). Bars and asterisks on the figure indicate significant differences between time-points for the CD8^high^ group. Additional significant comparisons are listed in text adjacent to the graph. (**C**) Percentage of CD8^+^ T cells expressing Granzyme B (GzmB^+^), in CD8^high^ (dark green) and CD8^low^ (black) patients with HCMV reactivation, and patients who did not experience HCMV reactivation (SN (blue), SP-NR (red)). Mean ± SEM are shown for each group. Statistically significant comparisons are indicated on the figure. The two patients with HCMV reactivation who lacked a T3 sample (n=1 LR, n=1 HR) are not shown in (**B**) or (**C**). Statistical significance was evaluated using mixed-effects models with Tukey’s multiple comparisons test (* p<0.05, ** p<0.01, *** p<0.001). SN, seronegative (n=11); SP-NR, seropositive no reactivation (n=5); LR, low-level reactivation (n=6); HR, high-level reactivation (n=13). T1 is prior to the detection of HCMV reactivation; T2, at the initial detection of HCMV reactivation; T3, the peak; T4, near the resolution of HCMV reactivation.

The CD8^high^ signature was more common in LR than HR, however there was no significant relationship with viral magnitude or days post-reactivation (**Table S6**). All CD8^low^ reactivators had received T cell depletion (TCD), whereas 6/10 CD8^high^ reactivators underwent TCD. The CD8^low^ group uniquely contained three patients who received bone-marrow grafts and one who experienced rejection (**Table S6**). These factors are associated with slower T cell recovery (Bosch et al., 2012; Schmidt-Hieber et al., 2010; Storek et al., 2001; Waller et al., 2019), thus the CD8^low^ signature may be reflective of conditioning and/or stem cell source.

We found that CD8^low^ patients had higher proportions of GzmB^+^ CD8^+^ T cells before the detection of HCMV reactivation (T1), compared to all other patient groups (**Figure 4C**). Thus, early enrichment of GzmB^+^ CD8^+^ T cells (above ∼74% of CD8^+^ T cells) may be a feature that heralds later development of HCMV reactivation and slower numeric CD8^+^ T cell recovery, particularly in TCD patients.

### MAIT cell frequencies at the initial detection of HCMV reactivation distinguish low-level and high-level reactivators

To further examine immune reconstitution differences between LR and HR patients, we performed SAM comparing the frequencies of all immune subsets examined between LR and HR. This revealed recovery of mucosal-associated invariant T (MAIT) cells and EM CD4^+^ T cells as key features distinguishing LR from HR patients (**Figure S5**).

We found an inverse relationship between the magnitude of HCMV DNAemia and MAIT cell frequency in patients with HCMV reactivation (**Figure 5**). Initial analysis focused on CD8^+^ MAIT cells (Vα7.2^+^CD161^high^ CD8^+^ T cells). CD8^+^ MAIT cell levels did not increase over time in any patient group, in line with reports of limited MAIT cell recovery in the first 1-2 years post-transplant (Bhattacharyya et al., 2018; Solders et al., 2017). However, MAIT cell absolute counts (**Figure 5A**) and percentages (**Figure 5B**) were consistently higher in LR than HR across the time-points. Importantly, MAIT cell levels were significantly higher in LR than HR at the initial detection of HCMV reactivation (T2) (**Figure 5A-B**), a time-point at which the magnitude of HCMV DNAemia (**Figure 1C**) and level of antigen exposure (HCMV AUC) were equivalent in LR and HR patients (**Figure S1C; Table S1**), and which was before (median 9 (26-7) days) any administration of pre-emptive antiviral therapy.

**Figure 5.**
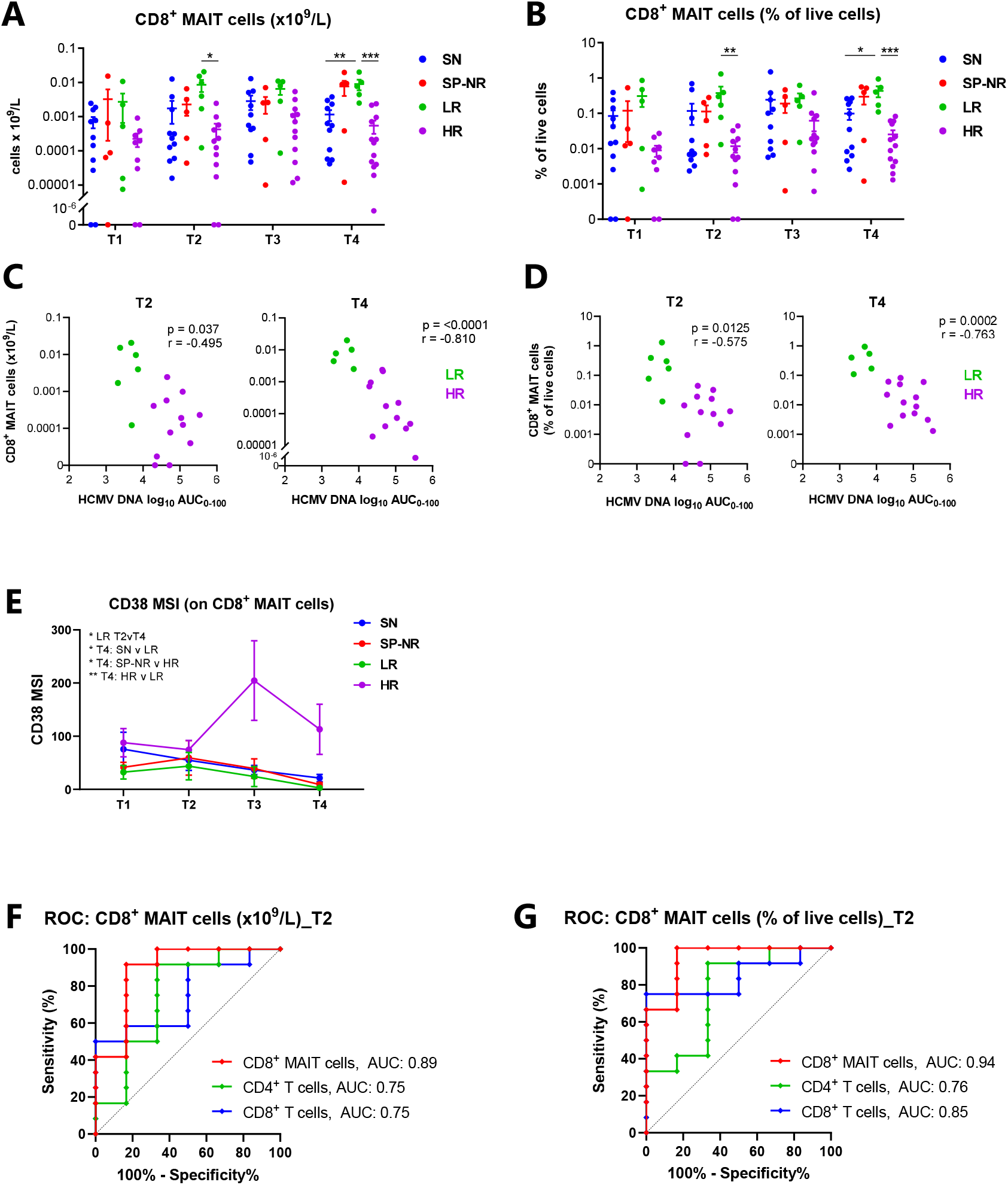
CD8^+^ MAIT cell frequencies at the initial detection of reactivation distinguish low-level and high-level HCMV reactivators. CD8^+^ MAIT cells were identified as Vα7.2^+^CD161^hi^CD8^+^ T cells. (**A**) Absolute counts (×10^9^/L blood) of CD8^+^ MAIT cells over the four study time-points. Mean ± SEM, mixed-effects model with Tukey’s multiple comparisons test (* p<0.05, ** p<0.01, *** p<0.001). (**B**) CD8^+^ MAIT cells as a percentage of total live cells. Mean ± SEM, mixed-effects model with Tukey’s multiple comparisons test (* p<0.05, ** p<0.01, *** p<0.001). (**C**) Spearman correlations at T2 (left) and T4 (right) between CD8^+^ MAIT cells (x10^9^/L) (from (A)) in patients with HCMV reactivation (LR (green), HR (purple)) and the log_10_ area under the curve (AUC) of HCMV DNA copies/mL over 0-100 days post-HSCT. At T2, LR (n=6) and HR (n=12). At T4, LR (n=5) and HR (n=13). (**D**) Spearman correlations at T2 (left) and T4 (right) between CD8^+^ MAIT cell percentages (from (B)) in patients with HCMV reactivation (LR (green), HR (purple)) and the log_10_ HCMV DNA AUC over 0-100 days post-transplant. (**E**) Median signal intensity (MSI) of CD38 on CD8^+^ MAIT cells. Mean ± SEM shown. Mixed-effects model with Tukey’s multiple comparisons test (* p<0.05, ** p<0.01); statistically significant comparisons are listed on the figure. (**F** and **G**) Receiver-operating characteristic (ROC) curves for the performance of CD8^+^ MAIT cells, CD8^+^ T cells and CD4^+^ T cells at T2 in discriminating LR and HR patients; (**F**) shows absolute counts (×10^9^/L), (**G**) shows cell subsets as percentages of total live cells. Optimal CD8^+^ MAIT cell cut-offs were (F) 0.001337 ×10^9^ cells/L (91.67% sensitivity, 83.33% specificity) and (G) 0.0605 % of live cells (100% sensitivity, 83.33% specificity). T1 is prior to the detection of HCMV reactivation; T2, at the initial detection of HCMV reactivation; T3, the peak; T4, near the resolution of HCMV reactivation. SN, HCMV-seronegative (n=11); SP-NR, seropositive no reactivation (n=5); LR, low-level HCMV reactivation (n=6); HR, high-level HCMV reactivation (n=13). AUC, area under the curve.

The median percentage of CD8^+^ MAIT cells at T2 was 40.3-fold higher in LR than HR patients (medians 0.235% and 0.006% of live cells, respectively). MAIT cell absolute counts and percentages were also significantly higher at T4 in LR compared to both HR and SN (**Figure 5A-B**). These findings held true when the total MAIT cell population (CD3^+^Vα7.2^+^CD161^high^) was examined without prior gating on CD8 (**Figure S6A-B**), and when MAIT frequencies were measured as a percentage of total CD3^+^ cells (**Figure S6C-D**).

CD8^+^ MAIT cell frequencies at T2 and T4 in patients with HCMV reactivation inversely correlated with the HCMV DNA AUC_0-100_ (**Figure 5C-D**) and peak HCMV copies/mL (**Figure S7**). Expression of the activation marker CD38 increased on CD8^+^ MAIT cells in HR, but not LR, at T3 and was significantly higher at T4 in HR compared to LR and SP-NR (**Figure 5E**).

We generated receiver-operating characteristic (ROC) curves to estimate the ability of CD8^+^ MAIT cell frequencies at T2 to discriminate patients who proceeded to develop low- or high-level HCMV reactivation (**Figure 5F-G**). Both CD8^+^ MAIT cell absolute counts (**Figure 5F**) and percentages of live cells (**Figure 5G**) demonstrated high AUCs of 0.89 (95%CI: 0.702-1.0; p=0.0087) and 0.94 (95%CI: 0.828-1.0; p=0.0027) respectively. CD8^+^ MAIT cells showed higher predictive ability than total CD8^+^ T cells or total CD4^+^ T cells (**Figure 5F-G**), which are parameters assessed in routine post-transplant monitoring. The highest combined sensitivity (100%) and specificity (83.33%) was achieved using a threshold value of 0.0605% CD8^+^ MAIT cells (% of live cells). 12/13 patients with <0.0605% CD8^+^ MAIT cells at T2 ultimately developed high-level HCMV reactivation (12 HR; 1 LR), while all (5/5) patients above this cut-off experienced low-level HCMV reactivation. Together, these data propose measurement of MAIT cell levels at the initial detection of HCMV DNAemia as a potential marker for predicting the subsequent magnitude of HCMV DNAemia and anticipating the need for antiviral therapy.

### Expansion of effector-memory CD4^+^ T cells in HSCT recipients with low-level HCMV reactivation

While EM CD4^+^ T cell levels were not significantly different between LR and HR before the detection of HCMV reactivation (T1), LR demonstrated faster expansion of EM CD4^+^ T cells over the course of reactivation, as reflected in absolute counts (**Figure 6A**) and percentages within the CD4^+^ T cell compartment (**Figure 6B**). Recovery of EM CD4^+^ T cell counts in HR patients remained low and was similar to non-reactivators (**Figure 6A**). Significantly higher EM CD4^+^ T cell percentages at T3 and T4 were seen in LR compared to HR (medians 75.0% vs. 41.6% at T3, and 76.9% vs. 42.5% at T4, for LR and HR respectively) (**Figure 6B**). The percentage of EM CD4^+^ T cells at T3 inversely correlated with the HCMV AUC_0-100_ (**Figure 6C**) and peak HCMV copy number (**Figure 6D**) in reactivators.

**Figure 6.**
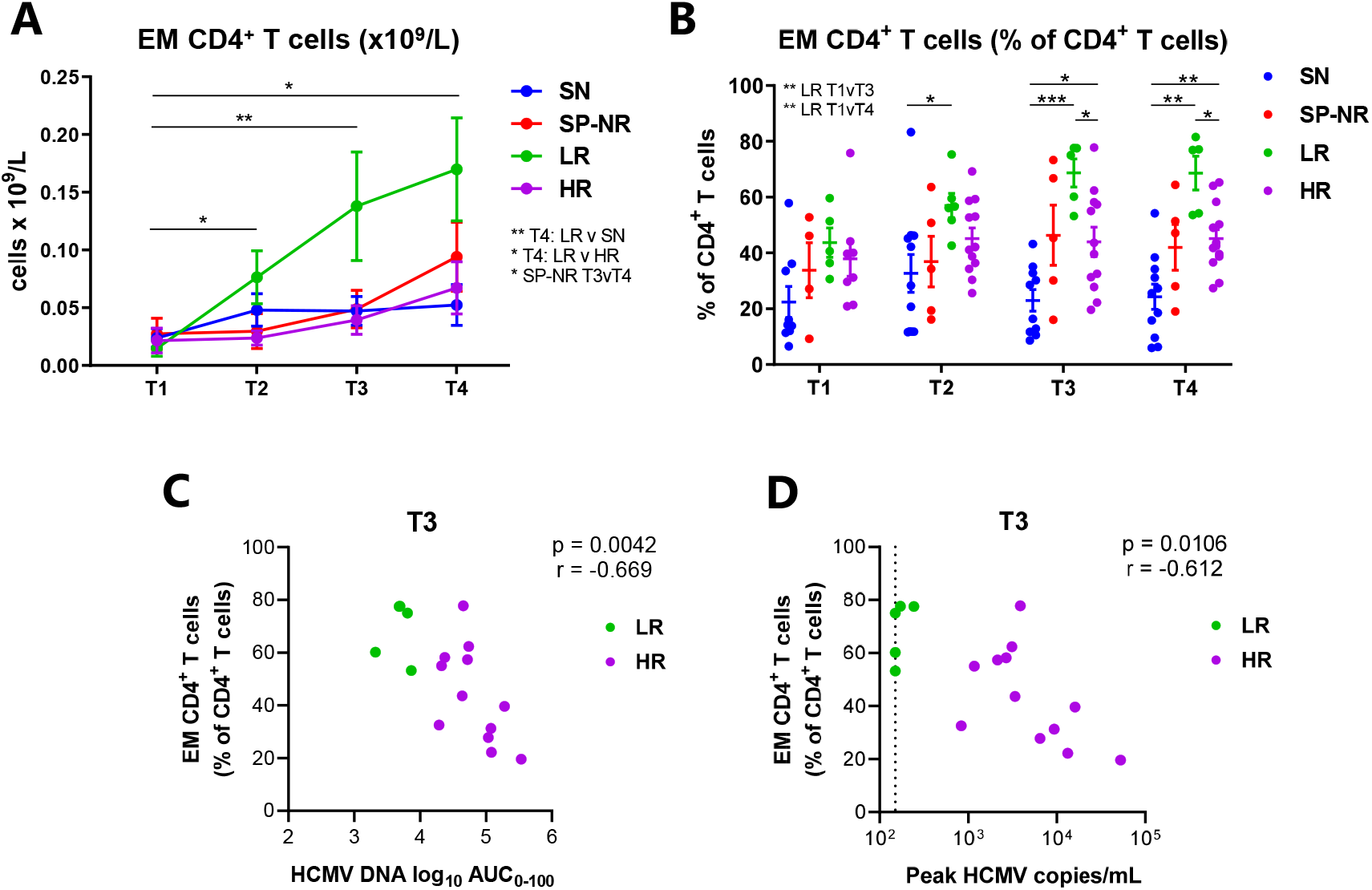
Expansion of effector-memory CD4^+^ T cells in HSCT recipients with low-level HCMV reactivation. (**A**) Absolute counts (x10^9^/L blood) of effector-memory (EM; CD45RA^-^CD45RO^+^CCR7^-^ CD27^-^) CD4^+^ T cells. Graph shows mean ± SEM. Mixed-effects model with Tukey’s multiple comparisons test (* p<0.05, ** p<0.01). Bars and asterisks indicate significant comparisons for LR between time-points. Additional significant comparisons are listed adjacent to the graph. (**B**) Percentage of CD4^+^ T cells with EM phenotype. Note: some green circle symbols (for LR patients) appear overlapping at T3 (77.6%, 77.5%) and T4 (54.2%, 53.6%; and 76.9%, 77.1%). Lines indicate mean ± SEM. Mixed-effects model with Tukey’s multiple comparisons test (* p<0.05, ** p<0.01, *** p<0.001). (**C**) Spearman correlation between log_10_ area under the curve (AUC) of HCMV DNA copies/mL over 0-100 days post-transplant, and EM CD4^+^ T cell percentages (of total CD4^+^ T cells) at T3 in HSCT recipients with HCMV reactivation (low-level (<250 copies/mL) reactivation (green; n=5 at T3); high-level (>830 copies/mL) reactivation (purple; n=12 at T3)). (**D**) Spearman correlation between peak HCMV copies/mL and EM CD4^+^ T cell percentages at the peak of HCMV reactivation (T3). The vertical black dotted line at 150 HCMV copies/mL plasma indicates the lower limit of quantitation of the HCMV DNA quantitative PCR assay. T1 is prior to the detection of HCMV reactivation; T2, at the initial detection of HCMV reactivation; T3, the peak; T4, near the resolution of HCMV reactivation. SN, HCMV-seronegative (blue; n=11); SP-NR, seropositive no reactivation (red; n=5); LR, low-level HCMV reactivation (green; n=6); HR, high-level HCMV reactivation (purple; n=13).

## DISCUSSION

This study details the evolution of changes in immune reconstitution at different phases of HCMV reactivation post-HSCT. We identified an inverse relationship between the magnitude of HCMV DNAemia and MAIT cell frequency in patients with HCMV reactivation, indicating MAIT cells may be a novel prognostic marker to determine which patients may benefit from early intervention. Stronger EM CD4^+^ T cell reconstitution was also evident in patients who experienced low-level HCMV reactivation, consistent with an important role in viral control.

A major obstacle in management of HCMV reactivation post-HSCT is potential antiviral drug-related toxicities from ganciclovir and foscarnet. Prophylaxis with safer agents such as Letermovir still carries a risk of viral resistance (Douglas et al., 2019), expense and the potential for delayed HCMV-specific T cell recovery (Zamora et al., 2021). Early discrimination of low-level and high-level reactivators could facilitate the tailoring of therapeutic decisions, as patients who develop self-limiting, low-level HCMV DNAemia may benefit from a shortened duration of prophylaxis or higher viral load threshold for pre-emptive therapy initiation.

The viral load at initial detection of HCMV reactivation has limited ability to predict which patients will develop clinically-significant infection (Camargo et al., 2018; Giménez et al., 2014). Building on work by us and others demonstrating the utility of mass cytometry to describe immune profiles associated with post-HSCT complications (Hartmann et al., 2019; Horowitz et al., 2015; Lakshmikanth et al., 2017; Li et al., 2019; McGuire et al., 2020; Stikvoort et al., 2017), here we stratified patients by HCMV-serostatus and the magnitude of peak HCMV DNAemia, directly analysing times aligning with HCMV viral load kinetics, including before and at initial detection of HCMV DNAemia.

A critical finding was that MAIT cell levels at the initial detection of HCMV DNAemia (cut-off 0.0605% of live cells) could effectively discriminate patients who eventually developed high-level HCMV DNAemia (>830 peak copies/mL) and required antiviral pharmacotherapy, versus those with low-level HCMV DNAemia (<250 peak copies/mL) who almost all cleared the infection spontaneously. MAIT cells are innate-like T cells with a semi-invariant T cell receptor (TCR) (Vα7.2) and recognise bacterial- and fungal-derived riboflavin metabolites presented on MHC class I-related molecule (MR1) (Kjer-Nielsen et al., 2012). MAIT cell levels were significantly lower in patients with high-level HCMV reactivation compared to low-level reactivation. To our knowledge, previous studies have not identified an association between MAIT cell recovery and infectious complications after allogeneic HSCT, including HCMV (Solders et al., 2017; Youssef et al., 2018). Reduced circulating MAIT cell frequencies have been observed in HCMV-seropositive healthy individuals (Patin et al., 2018) and depletion of circulating MAIT cells has been reported in several viral infections, linked to chronic immune activation (Barathan et al., 2016; Deschler et al., 2021; Fernandez et al., 2015; Flament et al., 2021; Hengst et al., 2016; Leeansyah et al., 2013; Paquin-Proulx et al., 2017; Parrot et al., 2020; Ussher et al., 2018; van Wilgenburg et al., 2016). Lower circulating MAIT cell frequencies have also been reported in severe GvHD (Bhattacharyya et al., 2018; Kawaguchi et al., 2018; Konuma et al., 2020; Stikvoort et al., 2017; van der Waart et al., 2012). In light of our findings, validation studies in a larger cohort will be important to evaluate the predictive ability of MAIT cell levels at the initial detection of HCMV reactivation in stratifying patients who subsequently develop low-level versus high-level HCMV reactivation.

Additional studies will also be needed to investigate the mechanisms underlying this difference in MAIT cell frequencies. Differences in thymic function, graft MAIT cell content and gut microbiota could be explored. Proliferation of graft-derived MAIT cells, requiring stimulation by both microbial MR1/TCR ligands and proinflammatory cytokines, is reported to contribute to early MAIT cell recovery post-HSCT (Bhattacharyya et al., 2018). Further, higher gut microbial diversity (Konuma et al., 2020) and abundance of Blautia spp. and Bifidobacterium longum (Bhattacharyya et al., 2018) are associated with better post-transplant MAIT cell recovery. The interaction between gut microbiota and immune cell reconstitution post-HSCT is becoming recognised (Fiorenza and Turtle, 2021; Ingham et al., 2019; Nguyen et al., 2021; Schluter et al., 2020), with low intestinal microbial diversity connected with increased mortality and inferior clinical outcomes (Peled et al., 2020). We found that MAIT cell levels inversely correlated with the HCMV DNA AUC_0-100_, a measure that is associated with increased risk of overall mortality (Hill et al., 2018; Stern et al., 2021). It is possible that poor MAIT cell recovery may be a biomarker of inferior overall immune reconstitution which predisposes to the development of high-level HCMV DNAemia.

Immunosuppressive therapies may also impact MAIT cell recovery. Post-HSCT cyclophosphamide has been associated with impaired MAIT cell recovery (Bhattacharyya et al., 2018), and cyclosporine A and sirolimus were found to impair MAIT cell proliferation *in vitro* (Solders et al., 2017). In our cohort, two patients in the HR group received conditioning with post-transplant cyclophosphamide and there was a greater proportion of patients with reduced intensity conditioning and grade II aGvHD in the HR group. Future studies with a larger number of patients are required to explore the impact of each of these variables further.

MAIT cells exhibit tissue-homing properties (Dusseaux et al., 2011), thus it might also be hypothesised that recruitment of blood MAIT cells to tissue sites of inflammation (van der Waart et al., 2012) may contribute to the lower frequency of circulating MAIT cells observed in HR. Against this, Youssef et al. (2018) were unable to detect TCR-Vα7.2^+^ lymphocytes in intestinal biopsies from paediatric HSCT recipients with aGvHD, HCMV reactivation or other viral infections. Examination of MAIT cells in HCMV-affected tissue biopsies in further studies represents an important avenue of research.

The functional role of MAIT cells in HCMV infection, if any, remains to be determined. MAIT cells can be activated in viral infections in a TCR-independent manner through inflammatory cytokines including IL-12 and IL-18 (Ussher et al., 2014; van Wilgenburg et al., 2016), and it is possible they may contribute to protective antiviral responses via the release of cytokines, such as IFN-γ and TNF, and cytotoxic granules (Loh et al., 2016; van Wilgenburg et al., 2016). Interestingly, HCMV encodes mechanisms to downregulate MR-1 *in vitro* (McSharry et al., 2020), suggesting the virus might benefit from evasion of TCR-dependent MAIT cell activation. Thus, investigation into the functional properties of MAIT cells in HSCT recipients with HCMV reactivation is warranted.

Our data also pointed to faster EM CD4^+^ T cell recovery over the course of HCMV reactivation as a key feature distinguishing low-level reactivators from high-level reactivators. This supports the importance of CD4^+^ T cell responses in the control of HCMV reactivation (Drylewicz et al., 2016; Einsele et al., 2002; Fabrizio et al., 2021; Gabanti et al., 2015; Gratama et al., 2008; Lilleri et al., 2012; Lim et al., 2020; Ljungman et al., 1993). Early HCMV-specific EM CD4^+^ T cell recovery post-HSCT has been associated with protection from HCMV reactivation (Pourgheysari et al., 2009). HCMV-specific CD4^+^ T cell recovery is needed for effective HCMV-specific CD8^+^ T cell responses and HCMV clearance (Foster et al., 2002; Muñoz-Cobo et al., 2012; Widmann et al., 2008).

No significant differences in immune composition were identified when directly comparing reactivators and non-reactivators before the detection of reactivation (T1; median day 19 post-HSCT). At this early time-point, there is likely to be substantial heterogeneity in immune recovery due to the impacts of transplant conditioning and differences in graft composition. Assessment of HCMV-specific CD8^+^ T cell functional profiles is a promising predictive method for distinguishing spontaneous controllers from those who progress to clinically-significant HCMV reactivation (Camargo et al., 2019; Tey et al., 2013). Limitations in cell numbers and blood volume in the pre-engraftment period often restrict analysis of PBMCs collected before the detection of HCMV reactivation, but here we utilised CyTOF to maximise the data extracted from such samples, which could in future help to refine immunological prediction of HCMV reactivation early post-transplant. We observed high proportions of GzmB-expressing CD8^+^ T cells prior to the detection of HCMV DNAemia in a subset of patients, which might reflect an early HCMV-specific T cell response to HCMV reactivation below the limit of detection.

Our observations including memory skewing, accumulation of late-differentiated T cells and inversion of the CD4:CD8 ratio are consistent with a prominent impact of HCMV reactivation on T cell reconstitution (Itzykson et al., 2015; Lugthart et al., 2014; Politikos et al., 2020; Suessmuth et al., 2015). The activated immune environment seen at the initial detection of HCMV DNAemia may reflect an early response to HCMV replication and/or it may provide an immune environment that favours HCMV reactivation (Reeves and Sinclair, 2013). Elevated frequencies of GzmB^+^ γδ T cells and NKG2C^+^CD57^+^ NK cells emerged with the progression of HCMV reactivation and have previously been implicated in HCMV control post-HSCT (Foley et al., 2012; Hammer et al., 2018; Kheav et al., 2014; Knight et al., 2010; Muñoz-Cobo et al., 2012; Ravens et al., 2017). A salient observation at T4 was significantly lower proportions of naïve CD8^+^ T cells and CD27^+^ T cells in reactivators compared to non-reactivators. Suessmuth et al. (2015) previously described a contraction of the naïve T cell compartment in HSCT patients with HCMV reactivation, and linked the clonal expansion of HCMV-specific EM CD8^+^ T cells to lower TCRβ diversity and defects in the TCRβ repertoire, suggestive of possible thymic impairment.

Although accelerated CD8^+^ T cell recovery is a canonical feature associated with HCMV reactivation post-HSCT (Drylewicz et al., 2016; Lugthart et al., 2014; Politikos et al., 2020; Suessmuth et al., 2015), we found that only a subgroup of reactivators developed a numerically elevated CD8^+^ T cell dominated immune profile. This ‘CD8^high^’ signature was seen in a majority of low-level reactivators, and indeed LR patients displayed the fastest overall CD8^+^ T cell recovery, suggesting this well-described observation is associated with viral control. The presence of HCMV-specific CD8^+^ T cells in seropositive recipients is reported to correlate with faster total CD8^+^ T cell reconstitution during the first year post-HSCT (Ogonek et al., 2017) and it will be informative to examine the antigen-specificity and function of the T cell subsets within this elevated quantitative profile. We previously found that development of an adaptive immune signature containing increased numbers of activated CD8^+^ T cells was associated with HCMV clearance in HSCT patients who received adoptive virus-specific T cell therapy for pharmaco-refractory HCMV reactivation (McGuire et al., 2020).

In conclusion, this study provides comprehensive insight into the evolution of immune reconstitution profiles at different phases of HCMV reactivation post-HSCT, with simultaneous examination of multiple innate and adaptive populations, encompassing both absolute cell counts and percentages, providing a nuanced perspective on patterns of immune reconstitution. Our findings suggest that MAIT cell levels at the initial detection of HCMV DNAemia could be further explored as a potential prognostic biomarker for anticipating the eventual magnitude of HCMV reactivation and requirement for antiviral therapy.

## Acknowledgements

This work was supported by a University of Sydney MCR-Biomed Connect Grant awarded to B.S. and a University of Sydney Kickstart Seed Grant awarded to H.M.M.. L.S. was supported by an Australian Government Research Training Program Scholarship. H.M.M. is currently supported by the International Society for the Advancement of Cytometry (ISAC) Marylou Ingram Scholars Program (2019-2023), and previously awarded an NHMRC post-doctoral fellowship [grant number GNT1037298]. The authors wish to thank Caryn van Vreden and all the support staff at Sydney Cytometry and the Ramaciotti Facility for Human Systems Biology, for their assistance with the mass cytometry studies, and Elissa Atkins and Angela Bayley from Westmead Hospital, for assistance with collection of patient data and samples.

## Author Contributions

B.S., A.A. conceived the study. B.S., L.S., A.A., E.B, H.M.M., S.A. designed the study. E.B, S.A., D.G. provided study materials. L.S. undertook the key experiments with assistance from H.M.M.. L.S., H.M.M., B.S., E.B, A.A., S.A., B.F. provided data analysis and interpretation, and all authors contributed to writing the manuscript and final approval of the submitted manuscript.

## METHODS

### Resource availability

#### Lead Contact

Further information and requests for resources and reagents should be directed to and will be fulfilled by the lead contact, Barry Slobedman.

#### Materials availability

This study did not generate new unique reagents.

#### Data availability

CyTOF data reported in this paper will be shared by the lead contact upon request. This paper does not report original code. Any additional information required to reanalyze the data reported in this paper is available from the lead contact upon request.

### Experimental model and subject details

#### Study subjects

This study included 35 adults who underwent peripheral blood (n=32) or bone marrow (n=3) allogeneic HSCT at Westmead Hospital (Sydney, Australia) between 2015 and 2017. Patient demographic and transplant characteristics are outlined in **Table 1**. Patients with cryopreserved PBMC samples available at ≥2 time-points in the retrospective study-design (see ‘Study design’ below) were included. All patients gave informed consent in accordance with the Declaration of Helsinki. This study was approved by the University of Sydney and Western Sydney Local Health District ethics committees.

Transplant conditioning regimens were myeloablative conditioning with cyclophosphamide-busulfan (n=11) or cyclophosphamide-total body irradiation (n=3); or reduced intensity conditioning with fludarabine-melphalan (n=16), fludarabine-cyclophosphamide (n=3), or fludarabine-cyclophosphamide with total body irradiation (one fraction) (n=2). Four patients with matched unrelated donors received Tocilizumab as a component of conditioning (n=3 SN, n=1 SP-NR). Twenty patients received *in vivo* T cell depletion with anti-thymocyte globulin (n=17) or alemtuzumab (n=3) (Table 1). Donor and/or recipient pre-transplant EBV serostatus was available for 34/35 patients; all (34/34) were donor and/or recipient EBV seropositive. GvHD prophylaxis was according to institutional protocols depending on the indication for transplant, donor source and conditioning regimen.

## Method Details

### Isolation of peripheral blood mononuclear cells

Peripheral blood was collected prospectively at weekly or greater intervals until day 100 post-HSCT, in EDTA vacutainers. Peripheral blood mononuclear cells (PBMCs) were isolated by Ficoll-Paque PLUS (GE Healthcare) density-gradient centrifugation, washed in DPBS (without Ca or Mg; Lonza), and cryopreserved in freezing media (RPMI-1640 [with L-glutamine; Lonza] supplemented with 20% (v/v) foetal bovine serum [FBS; Sigma Aldrich] and 10% (v/v) dimethyl sulfoxide). Cells were stored at - 80°C for up to 7 days, then transferred to vapour-phase liquid nitrogen until use.

### Virological monitoring

Patients underwent weekly monitoring for HCMV and Epstein-Barr virus (EBV) in the first 100 days post-transplant. HCMV DNA load was measured in EDTA plasma by quantitative PCR (COBAS® AmpliPrep/COBAS® TaqMan® CMV Test; Roche). HCMV reactivation (also referred to as DNAemia) was defined as any detectable HCMV DNA in plasma. The lower limit of quantitation (LLQ) was 150 HCMV DNA copies/mL. The conversion to international units (IU) is 1 copy = 0.91 IU. EBV reactivation was defined as detection of EBV DNA at any level of quantitation using the EBV ELITe MGB® Kit (Elitech, Puteaux, France); the linear range of the assay was 150 to 10,000,000 copies/mL.

For graphical and calculation purposes, detectable HCMV DNA levels below the LLQ (<150 copies/mL) were assigned the value of 150 copies/mL. ‘Not detected’ readings (undetectable viral load) were assigned the value of 0 copies/mL. One HCMV-seronegative (D-/R-) patient had detection of HCMV DNA below the LLQ at a single time-point; this was considered a false positive and the patient was not excluded.

The log_10_ area under the curve (AUC) of HCMV DNA copies/mL plasma was calculated in GraphPad Prism version 8.2.1 (GraphPad Software, LLC) using the trapezoid rule. The duration of HCMV reactivation was calculated by first day minus last day of detected HCMV DNAemia, with any non-consecutive HCMV DNAemia episodes, including those extending beyond the first 100 days post-transplant, added together.

### Antiviral therapy for HCMV reactivation

HCMV-seropositive recipients received ganciclovir or valganciclovir prophylaxis from day -10 until day -1 pre-HSCT. Post-transplant HCMV reactivation was managed with pre-emptive antiviral therapy based on quantitative HCMV DNAemia measured weekly on peripheral blood plasma. Post-transplant pre-emptive antiviral therapy with ganciclovir and/or foscarnet was administered to 14/19 patients with HCMV reactivation. Antiviral pharmacotherapy was commenced when 2 consecutive HCMV PCR results showed increasing copy numbers, if a single result was greater than 1000 copies/mL or according to physician discretion if a HCMV viraemic patient was considered to be at high risk of uncontrolled HCMV replication. Ganciclovir 5 mg/kg IV 2 times daily (or dose equivalent of oral valganciclovir) was the first line agent and Foscarnet 60 mg/kg 3 times daily was used if (val)ganciclovir was contraindicated. A primary course of antiviral pharmacotherapy lasted 14 days, after which further treatment depended on response.

The point of pre-emptive therapy initiation for each patient is indicated on the HCMV DNAemia graphs in **Figure 1A** by the shift from the solid line (before therapy) to dotted line (after initiation of therapy). Of the 14 patients who received pre-emptive antiviral therapy for HCMV reactivation, 8/14 patients had received pre-emptive therapy at or before T3, and 14/14 patients had received pre-emptive therapy by T4. No patient received pre-emptive antiviral therapy at or before T2.

### Study design

The patients were retrospectively divided into four groups according to pre-transplant HCMV serostatus and magnitude of post-transplant HCMV reactivation (**Figure 1A**). The ‘*Seronegative*’ (SN; n=11) group were HCMV-seronegative recipients with seronegative donors (D-/R-), and had no detected HCMV DNAemia. The ‘*Seropositive No Reactivation*’ (SP-NR; n=5) group were HCMV-seropositive recipient (R+) and/or donor (D+) patients with no documented HCMV reactivation in the first 100 days post-HSCT. ‘*Low-level reactivators*’ (LR; n=6) developed HCMV reactivation to <250 peak copies/mL. ‘*High-level reactivators*’ (HR; n=13) developed HCMV reactivation to >830 peak copies/mL. No patients had peak HCMV titres between 250-830 copies/mL.

Mass cytometry was performed on PBMC samples from four time-points retrospectively selected in relation to the course of HCMV DNAemia (**Figure 1B**): (T1) prior to detection of HCMV DNAemia, (T2) at the initial detection of DNAemia, (T3) at the peak of HCMV DNAemia, and (T4) near the resolution of HCMV DNAemia (**Table S2**). Samples from matched days post-transplant were analysed from HSCT recipients who did not experience HCMV reactivation (SN and SP-NR) (**Figure 1D**).

A total of 130 PBMC samples were analysed. 9/35 patients (n=5 HR, n=3 LR, n=1 SN) did not have samples available at all four time-points. Of these patients, 8/9 were missing a sample from a single time-point, and 1/9 was missing samples from two time-points. There was a median of 19 (5-49) days between consecutive time-point samples analysed from an individual patient.

If cryopreserved PBMCs were not available from the precise days of T2, T3, or T4 time-points, the most proximate samples (where available) were used. For T2, if a PBMC sample from the day of first detected HCMV DNAemia was not available, the immediately succeeding available sample was used. If the precise T3 sample was not available, the most proximate sample immediately preceding the HCMV DNAemia peak was used. For T4, the sample at or most proximate to HCMV DNAemia resolution (undetectable viral load) in first 100 days post-transplant was analysed. At T4, with the exception of 3 HR patients, HCMV DNAemia was undetectable (6/13 HR and 6/6 LR) or had declined below the LLQ (4/13 HR). For patients with unresolved HCMV DNAemia in the first 100 days post-HSCT (n=7), T4 samples corresponded to suppression of HCMV DNAemia below the LLQ (n=4) or otherwise matched days post-HSCT to the other HR patients (n=3). The magnitude of HCMV DNAemia (in copies/mL plasma) at T4 in the patients with detectable HCMV DNAemia at T4 had decreased from peak levels by a median of 97.7% (range 83.2-98.8%).

### Mass cytometry staining, acquisition and analysis

#### Cell staining

PBMCs were stained for mass cytometry in eight batches using a panel of 36 metal-conjugated antibodies (**Table S3**). All samples from an individual patient were included in the same staining batch. Each batch contained a representation of patient groups (SN, SP-NR, LR, HR) and a batch control PBMC sample from a healthy donor. All antibodies were validated, pre-titered and supplied in per-test amounts by the Ramaciotti Facility for Human Systems Biology Mass Cytometry Reagent Bank. Reagent Bank antibodies were purchased either pre-conjugated from Fluidigm, or in a purified, carrier-free unlabelled format from other suppliers. Unlabelled antibodies were conjugated with the indicated metal isotope using the Maxpar Antibody Labelling Kit (Fluidigm, South San Francisco, CA) according to the manufacturer’s protocol. All antibodies were titrated before inclusion in the panel.

Briefly, cryopreserved PBMCs were quickly thawed in a 37°C waterbath and washed with warm RPMI-1640 (Lonza) supplemented with 10% (v/v) FBS (Sigma) and 1:10000 (v/v) Pierce Universal Cell Nuclease (Thermo Fisher Scientific). Cells were then washed with warm RPMI-1640 containing 10% FBS and counted using a haemocytometer and trypan blue. After one wash in serum-free RPMI-1640, up to 3.0 ×10^6^ viable cells per sample were retained for subsequent staining steps. For live/dead discrimination, cells were incubated with 1.25 μM Cell-ID™ cisplatin (Fluidigm) in serum-free RPMI-1640 for 3 min at room temperature (RT), followed by immediate quenching with RPMI-1640 containing 10% FBS. Cells were then washed once with FACS buffer (DPBS containing 1% FBS and 0.01 M EDTA) then incubated in a 50 μL surface antibody cocktail (prepared in FACS buffer) for 30 min at 4°C. The antibody cocktail mastermix was freshly prepared and centrifuged at 12,000 × g for 4 min through a 0.1 μm filter immediately before use. Following two washes with FACS buffer, cells were fixed and permeabilised by incubation in 1X FoxP3 Fixation/Permeabilisation buffer (eBioscience) for 45 min at 4°C, then washed twice with 1X Permeabilisation buffer (eBioscience). Cells were stained with a 50 μL intracellular antibody cocktail (mastermix prepared in 1X Permeabilisation buffer and centrifuged through a 0.1 μm filter immediately before use) for 45 min at 4°C. The cells were subsequently washed with 1X Permeabilisation buffer, then FACS buffer. Finally, cells were fixed in a 4% paraformaldehyde solution (prepared in PBS) containing 0.125 μM Cell-ID™ Iridium-191/193 nucleic acid intercalator (Fluidigm) for 20 min RT initially, then stored at 4°C overnight or for up to one week prior to acquisition.

#### Acquisition

On the day of acquisition, cells were washed once in FACS buffer, once in DPBS and then twice in MilliQ water. The pellet was resuspended at 0.8 ×10^6^ cells/mL in 1:10 (v/v) EQ Four Element Calibration Beads (Fluidigm) in MilliQ water, and filtered through a 0.35 μm nylon cell strainer snap-cap (Falcon) immediately prior to acquisition. Samples were acquired on CyTOF 2 Helios upgraded mass cytometer (Fluidigm) with an acquisition rate of up to 400 events/s and flow rate of 30 μL/min. A median of 311867 events were acquired per sample (n=130). FCS files were normalised on the CyTOF Software (version 6.7.1014) using the signal intensity of the EQ beads acquired within each sample.

#### Analysis

Cell populations were identified via manual gating in FlowJo 10.0.7 (Tree Star, Inc.). The gating strategy for major subsets is outlined in **Figure S2**. A list of all cell subsets analysed can be found in **Table S4** and **Table S5**. Analysis of cell subset percentages included both percent of live cells (% live) and/or percent of parent subset. Percentages of live cells (% live) were derived from the “live cells” gate indicated in **Figure S2**. Apart from major cell subsets, data were treated as missing if there were too few cells in the parent gate to allow for accurate gating of subpopulation(s). In these cases, the absolute counts were returned as zero.

Absolute cell counts (x10^9^/L blood) for each subset were calculated with reference to absolute monocyte (Mo) or lymphocyte (Ly) counts measured on a Sysmex 1800 full blood analyser on the day of PBMC sample collection. Briefly, the frequency of each cell subset as a percentage of total live cells was divided by either the sum of monocyte populations (classical, intermediate, non-classical) or the sum of lymphocyte populations (B cells, CD3^+^ cells, NK cells, mDC and pDC) as percentages of total live cells, and then multiplied by the absolute Mo or Ly count (x10^9^/L) (McGuire et al., 2020). Readings of ‘ND’ on the full blood analyser were assigned the value 0.5 ×10^8^ cells/L (half of the lowest quantified concentration for Ly and Mo). White blood cell (WBC) counts (x10^9^/L) were derived directly from the full blood analyser.

### Statistical Analysis

All statistical tests were performed using GraphPad Prism Software version 8.2.1 (GraphPad Software, LLC) unless indicated otherwise. To compare patient characteristics between groups, the Fisher’s exact test (categorical variables) or Mann-Whitney test (continuous variables) were used as appropriate. All tests were two-sided and comparisons with p<0.05 were considered significant. Two-class unpaired significance analysis of microarrays (SAM) (Tusher et al., 2001) in MultiExperiment Viewer 4.9.0 (TM4) (Saeed et al., 2003) was used to compare cell-subset frequencies between pairs of patient groups. Cell proportions (88 subsets) and absolute counts (77 subsets) were analysed independently for each time-point, using 100 permutations with delta adjusted to yield a false positive rate of zero. Heat map columns represent individual patients and are preserved in the same order across heat maps, except where samples were unavailable per time-point.

To further examine significant cell subsets as identified by SAM, two way mixed-effects models with the Geisser-Greenhouse correction followed by Tukey’s multiple comparisons test were used to enable comparison of cell subset frequencies both (a) between the four patient groups at each of four time-points, and (b) between the four time-points for each patient group. The mixed effects model is similar to repeated measures ANOVA, but allows for missing values. The mixed model implemented in GraphPad Prism uses a compound symmetry covariance matrix and is fit using restricted maximum likelihood (REML). The absolute counts of immune cell subsets were observed to have positively skewed distributions, thus data were log_10_-transformed to satisfy the assumption of normality prior to fitting the mixed effects model. Residual and QQ normality plots were visualised to confirm suitability of the model on the log-normal distribution. All absolute cell count graphs display untransformed data, with significance evaluated using log-transformed values. Correlations were evaluated using two-tailed Spearman correlations. Receiver-operating characteristic (ROC) curves were generated in GraphPad Prism 8.3.1. Data stated in the main text is the median with range in brackets.

