## Supplemental figures and tables for "Immune Cell Profiling Reveals MAIT and Effector Memory CD4+ T Cell Recovery Link to Control of Cytomegalovirus Reactivation after Stem Cell Transplant"

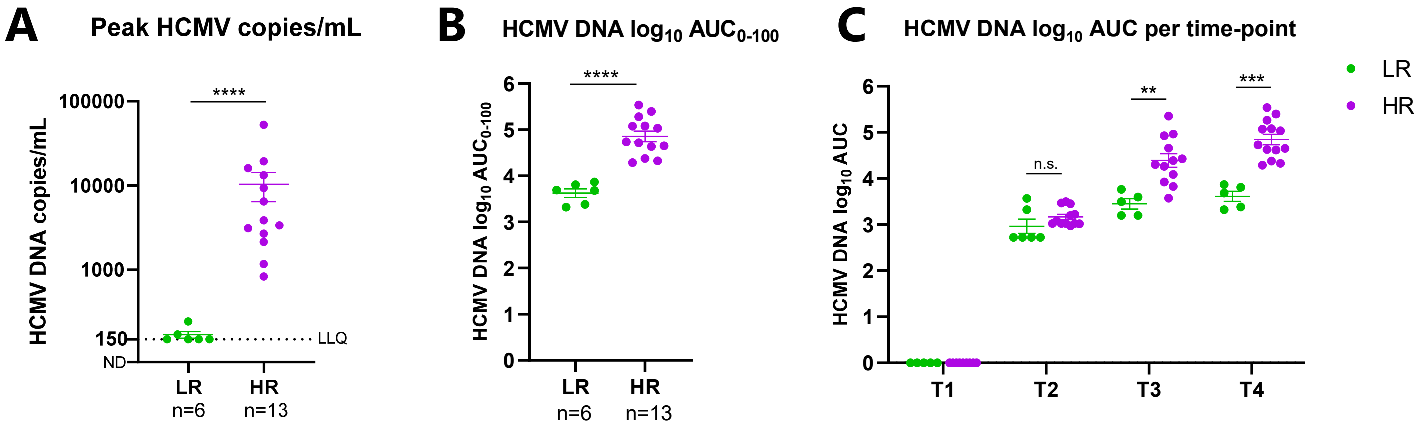


**<u>Supplemental Figure 1.</u> Magnitude of HCMV reactivation in HSCT recipients.**
(**A**) Peak HCMV copies/mL plasma in HSCT recipients with low-level (LR; green) or high-level (HR; purple) HCMV reactivation. The LLQ (lower limit of quantitation; 150 copies/mL) of the HCMV DNA quantitative PCR assay is indicated by the horizontal black dotted line. ND, not detected. Statistical significance between LR and HR evaluated using a Mann-Whitney U test (**** $p<0.0001$). (**B**) The HCMV DNA $log_{10}$ area under the curve (AUC) in the first 100 days post-HSCT ($AUC_{0-100}$) in LR and HR patients. Statistical significance between LR and HR evaluated using a Mann-Whitney U test (**** $p<0.0001$). (**C**) The HCMV DNA $log_{10}$ AUC in LR and HR patients at each time-point (T) analysed by mass cytometry. Statistical significance between LR and HR was evaluated per time-point using a Mann-Whitney U test (** $p<0.01$, *** $p<0.001$). Graphs show mean ± SEM. n.s., not significant.

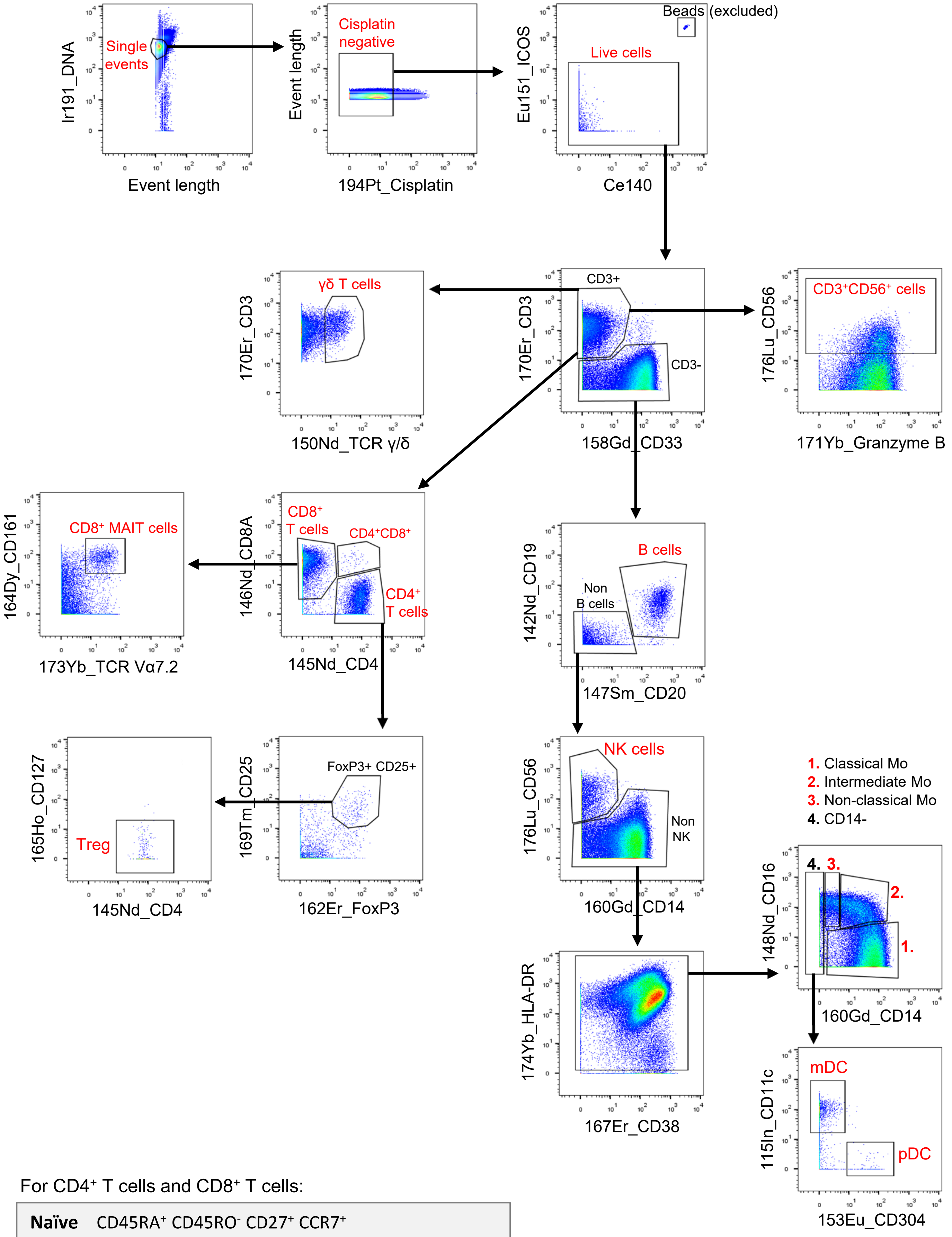


For CD4+ T cells and CD8+ T cells:

| | |
|---|---|
| **Naïve** | CD45RA+ CD45RO- CD27+ CCR7+ |
| **Central memory (CM)** | CD45RA- CD45RO+ CD27+ CCR7+ |
| **Effector memory (EM)** | CD45RA- CD45RO+ CD27- CCR7- |
| **EMRA** | CD45RA+ CD45RO- CD27- CCR7- |

**Supplemental Figure 2.** **Mass cytometry gating strategy for major subsets.**

**<u>Supplemental Figure 2.</u> Mass cytometry gating strategy for major subsets.**

Single events were distinguished from doublets and debris by DNA intercalator staining and event length. Cisplatin-negative (live) cells were then selected and EQ beads ($Eu151^{+}Ce140^{+}$) excluded prior to identification of major PBMC populations, as per the gating strategy shown. The 'live cells' gate indicated was the gate from which percentages of immune cell subsets (out of total live single cells) were derived. The phenotype used for identification of naïve and memory T cell subsets is shown (grey box). Further gating on cell subpopulations as listed in Supplemental Methods was conducted. All gating was performed in FlowJo software version 10.0.7 (Tree Star, Inc.). Mo, monocyte.

### A. Proportions

**T2**

- CD86$^+$ (% of classical mo)
- CD86$^+$ (% of intermediate mo)
- CD86$^+$ (% of mDCs)
- CD86$^+$ (% of non-classical mo)
- CD38$^+$ (% of CD8$^+$ MAIT cells)
- HLA-DR$^+$CD38$^+$ (% of CD8$^+$ T cells)
- HLA-DR$^+$CD38$^+$ (% of EM CD8$^+$ T cells)
- ICOS$^+$ (% of CD4$^+$ T cells)
- ICOS$^+$ HLA-DR$^+$CD38$^+$ (% of CD4$^+$ T cells)

**T3**

- HLA-DR$^+$CD38$^+$ (% of EM CD8$^+$ T cells)
- ICOS$^+$ (% of CD4$^+$ T cells)
- ICOS$^+$ HLA-DR$^+$CD38$^+$ (% of CD4$^+$ T cells)
- CM (% of CD8$^+$ T cells)
- CD38$^+$ (% of non-classical mo)
- CD38$^+$GzmB$^+$ (% of CM CD8$^+$ T cells)
- CD57$^+$GzmB$^+$ (% of CD8$^+$ T cells)
- HLA-DR$^+$CD38$^+$ EM (% of CD8$^+$ T cells)
- EM (% of CD8$^+$ T cells)
- GzmB$^+$ (% of CD8$^+$ T cells)
- CD57$^+$ (% of CD4$^+$ T cells)
- GzmB$^+$ (% of CD4$^+$ T cells)
- GzmB$^+$ (% of γδ-T cells)

**T4**

- CD38$^+$ (% of non-classical mo)
- CD38$^+$GzmB$^+$ (% of CM CD8$^+$ T cells)
- CD57$^+$GzmB$^+$ (% of CD8$^+$ T cells)
- HLA-DR$^+$CD38$^+$ EM (% of CD8$^+$ T cells)
- EM (% of CD8$^+$ T cells)
- GzmB$^+$ (% of CD8$^+$ T cells)
- CD57$^+$ (% of CD4$^+$ T cells)
- GzmB$^+$ (% of CD4$^+$ T cells)
- GzmB$^+$ (% of γδ-T cells)
- CD57$^+$NKG2C$^+$ (% of NK cells)
- CD57$^+$ (% of CD8$^+$ T cells)
- CD57$^+$ CD8$^+$ T cells (% of live cells)
- CD57$^+$CD38$^+$ (% of CD8$^+$ T cells)
- CD8$^+$ T cells (% of live cells)
- CD45RA$^+$GzmB$^+$CD27$^-$ (% of γδ-T cells)
- ↓ CD27$^+$ (% of γδ-T cells)
- ↓ CM (% of CD4$^+$ T cells)
- ↓ CD27$^+$ (% of CD4$^+$ T cells)
- ↓ CD27$^+$ (% of CD8$^+$ T cells)
- ↓ Naïve (% of CD8$^+$ T cells)

**T2 T3 T4**

- CD38+ (% of CD4+ T cells)
- PD-1+ (% of CD4+ T cells)
- HLA-DR+CD38+ (% of CD4+ T cells)
- EM (% of CD4+ T cells)
- HLA-DR+CD38+ EM (% of CD4+ T cells)
- HLA-DR+CD38+ (% of EM CD4+ T cells)

Key
**Monocyte/DC**
**NK cell**
**CD8$^+$ T cell**
**CD4$^+$ T cell**
**γδ-T cell**

### B. Absolute counts

**T2**

- CD27$^+$ CD4$^+$ T cells
- CM CD4$^+$ T cells

**T3**

- CD57$^+$PD-1$^+$CD38$^+$ CD4$^+$ T cells
- CD45RA$^+$GzmB$^+$CD27$^-$ γδ-T cells
- PD-1$^+$ CD8$^+$ T cells
- CD8$^+$ T cells
- CD38$^+$GzmB$^+$ CM CD8$^+$ T cells
- CD57$^+$ CD8$^+$ T cells
- HLA-DR$^+$CD38$^+$ CD8$^+$ T cells
- EM CD8$^+$ T cells
- CD57$^+$CD38$^+$ CD8$^+$ T cells
- HLA-DR$^+$CD38$^+$ EM CD8$^+$ T cells
- PD-1$^+$ EM CD8$^+$ T cells
- Tim-3$^+$ CD8$^+$ T cells
- CD161$^{lo/-}$ Vα7.2$^+$ CD8$^+$ T cells
- CM CD8$^+$ T cells
- GzmB$^+$ CD8$^+$ T cells
- CD57$^+$GzmB$^+$ CD8$^+$ T cells
- γδ-T cells
- CD45RA$^+$ γδ-T cells
- GzmB$^+$ γδ-T cells
- GzmB$^+$ CD4$^+$ T cells
- CD3$^+$CD56$^+$ cells
- CD57$^+$GzmB$^+$ CD3$^+$CD56$^+$ cells

**T4**

- Tim-3$^+$ CD8$^+$ T cells
- CD161$^{lo/-}$ Vα7.2$^+$ CD8$^+$ T cells
- CM CD8$^+$ T cells
- GzmB$^+$ CD8$^+$ T cells
- CD57$^+$GzmB$^+$ CD8$^+$ T cells
- γδ-T cells
- CD45RA$^+$ γδ-T cells
- GzmB$^+$ γδ-T cells
- GzmB$^+$ CD4$^+$ T cells
- CD3$^+$CD56$^+$ cells
- CD57$^+$GzmB$^+$ CD3$^+$CD56$^+$ cells
- NKG2C$^+$ CD8$^+$ T cells
- CD57$^+$NKG2C$^+$ NK cells
- CD16$^+$ γδ-T cells

### C. Percents and Counts

**T3**

- CD38$^+$GzmB$^+$ CM CD8$^+$ T cells
- EM CD8$^+$ T cells
- HLA-DR$^+$CD38$^+$ EM CD8$^+$ T cells
- CM CD8$^+$ T cells
- GzmB$^+$ CD8$^+$ T cells
- CD57$^+$GzmB$^+$ CD8$^+$ T cells
- GzmB$^+$ γδ-T cells

**T4**

- CM CD8$^+$ T cells
- GzmB$^+$ CD8$^+$ T cells
- CD57$^+$GzmB$^+$ CD8$^+$ T cells
- GzmB$^+$ γδ-T cells
- CD57$^+$NKG2C$^+$ NK cells

**Supplemental Figure 3.** Schematic representation of all cell subsets shown on heat maps in Figure 3.

**<u>Supplemental Figure 3.</u> Schematic representation of all cell subsets shown on heat maps in Figure 3.** The heat map rows in Figure 3 show cell subsets which differed significantly in percentage (Figure 3A) or absolute count (Figure 3B) in reactivators (LR and HR) compared to non-reactivators (SP-NR and SN), as determined by significance analysis of microarrays (SAM). (**A**) Visual representation of all cell subsets shown on Figure 3A heat maps, with text coloured by cell type. Coloured vertical lines adjacent to cell subsets indicate the time-point(s) at which the cell subsets were significantly higher in HCMV reactivation patients (LR, HR), compared to patients without reactivation (SN, SP-NR). The downward arrows at T4 are adjacent to cell subsets that were significantly lower in reactivators compared to non-reactivators. (**B**) Visual representation of all cell subsets displayed on Figure 3B heat maps. (**C**) Cell subsets which were significantly higher in both percentage (Figure 3A) and absolute count (Figure 3B) in reactivators compared to non-reactivators. T1 is prior to the detection of HCMV reactivation; T2, at the initial detection of HCMV reactivation; T3, the peak; T4, near the resolution of HCMV reactivation. SN, HCMV seronegative (n=11); SP-NR, seropositive no reactivation (n=5); LR, low-level HCMV reactivation (n=6); HR, high-level HCMV reactivation (n=13).

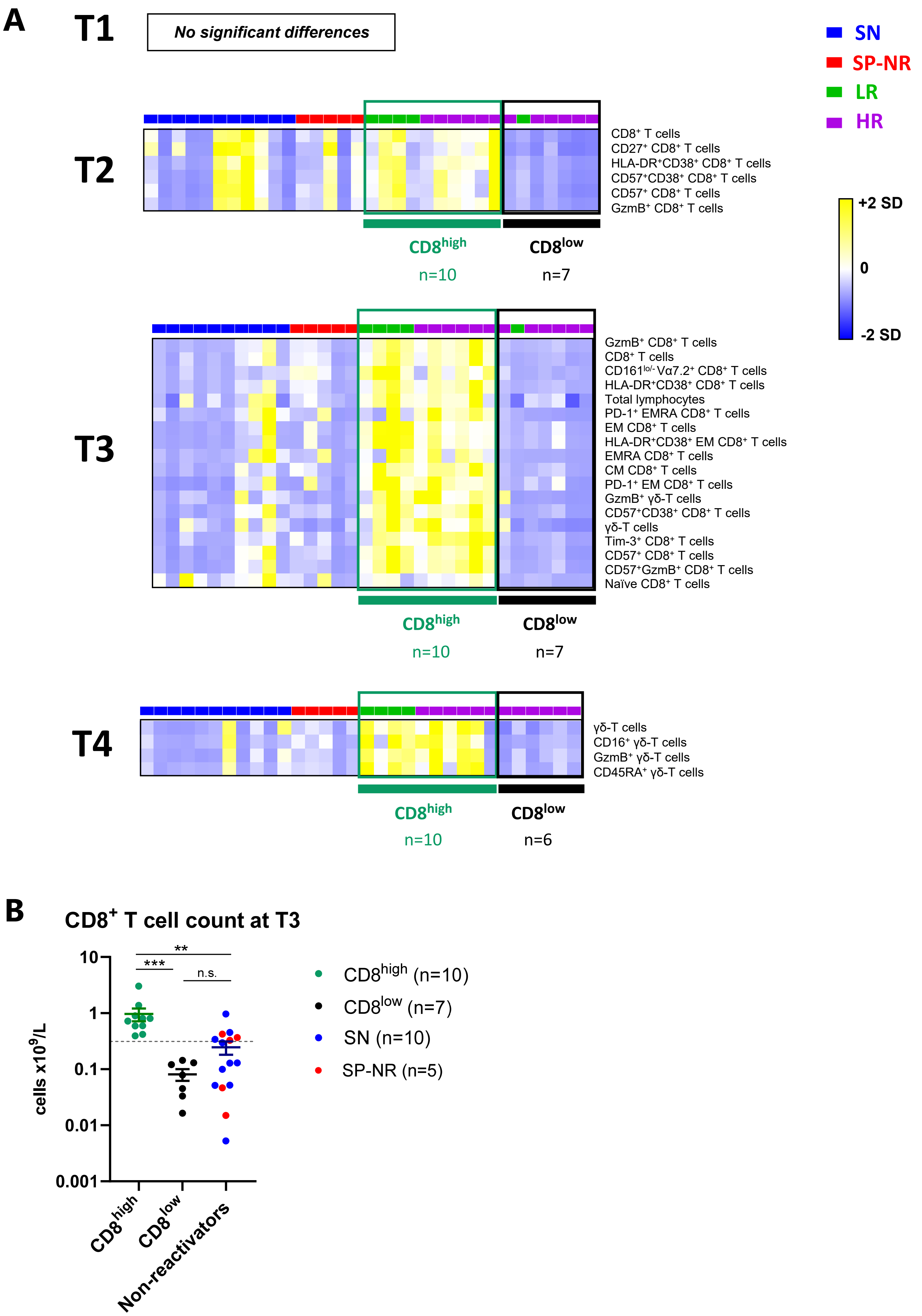


**Supplemental Figure 4.** Elevated T cell signature in CD8$^{high}$ reactivators compared to CD8$^{low}$ reactivators.

**<u>Supplemental Figure 4.</u> Elevated T cell signature in CD8$^{high}$ reactivators compared to CD8$^{low}$ reactivators.**

HSCT recipients with HCMV reactivation were divided into 'CD8$^{high}$' (n=10) and 'CD8$^{low}$' (n=7) groups after detection of a CD8$^{+}$ T cell dominated immune signature at T3 (peak of HCMV DNAemia) in Figure 3B in a subgroup of reactivators (termed 'CD8$^{high}$'). (**A**) Absolute counts of all (77) immune subsets were directly compared between CD8$^{high}$ reactivators (dark green) and CD8$^{low}$ reactivators (black) via two-class unpaired significance analysis of microarrays (SAM). Heat map rows display cell subsets that were significantly different in absolute count (x$10^9$/L blood) between CD8$^{high}$ and CD8$^{low}$ reactivators. As the CD8$^{high}$ and CD8$^{low}$ groups were defined at T3 (in Figure 3B), the two patients with HCMV reactivation who lacked a T3 sample (n=1 LR, n=1 HR) were not included in this analysis. Heat maps are coloured by the Z-score normalised per row. Each column represents a patient. SN (HCMV seronegative; blue) and SP-NR (seropositive no reactivation; red) patients are included on the heat maps for visual comparison. T1 is prior to the detection of HCMV reactivation; T2, at the initial detection of HCMV reactivation; T3, the peak; T4, near the resolution of HCMV reactivation. LR, low-level HCMV reactivation (light green); HR, high-level HCMV reactivation (purple); SD, standard deviation. (**B**) Absolute CD8$^{+}$ T cell counts (x$10^9$/L blood) at T3 (the peak of HCMV DNAemia) in CD8$^{high}$ reactivators (dark green), CD8$^{low}$ reactivators (black), and non-reactivators (SN (blue) and SP-NR (red)) are shown. The black horizontal dotted line indicates the median CD8$^{+}$ T cell count at T3 (0.3116 x$10^9$/L). Mean ± SEM is indicated. Statistical differences were determined by Kruskal-Wallis test with Dunn's multiple comparisons test (** $p<0.01$, *** $p<0.001$). n.s., not significant.

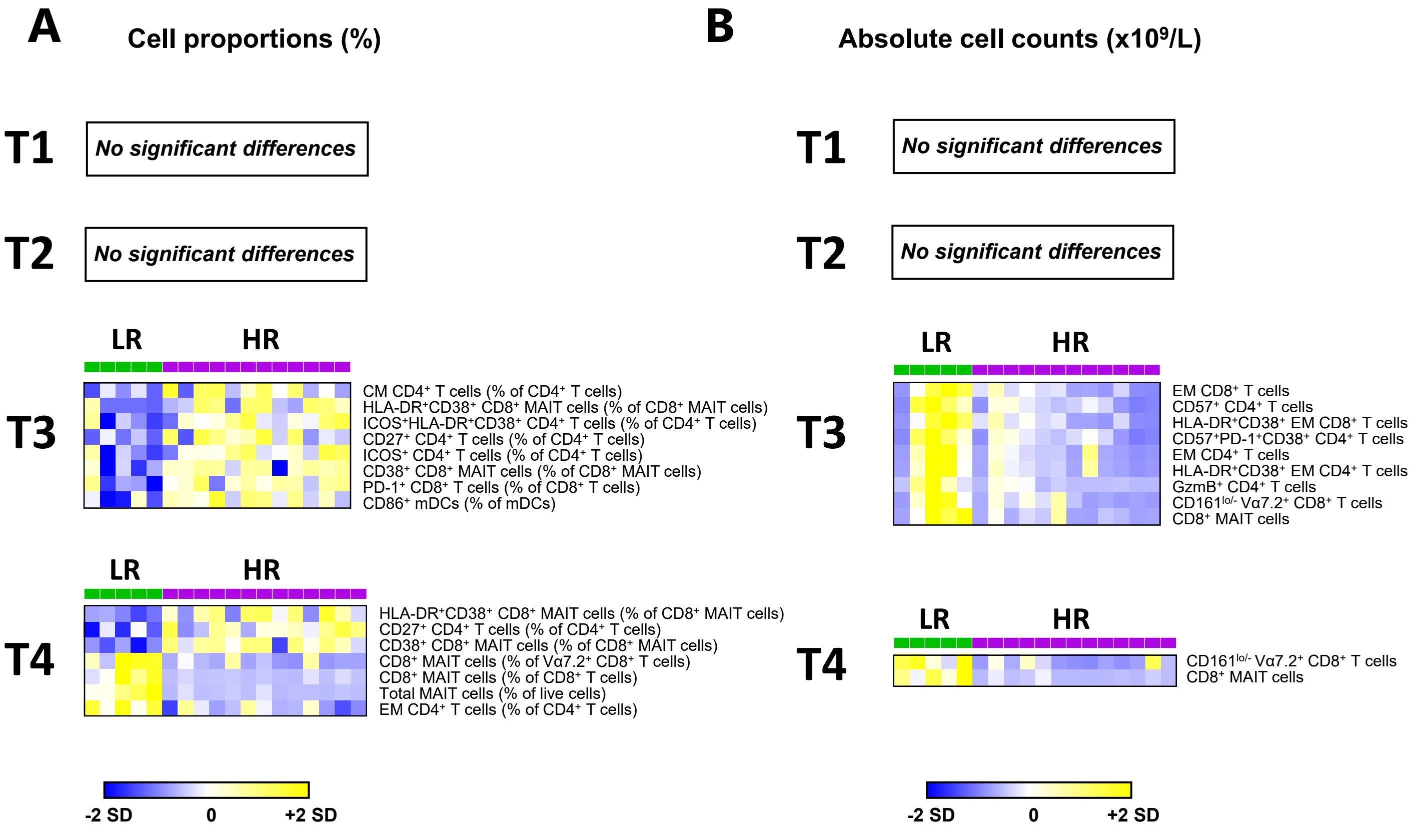


**<u>Supplemental Figure 5.</u> Immune cell subset differences in HSCT recipients with low-level HCMV reactivation versus high-level HCMV reactivation.**

Heat map rows display (**A**) percentages or (**B**) absolute counts (x10$^9$/L) of immune cell subsets that were significantly different between LR (green) and HR (purple) patients, as determined by two-class unpaired significance analysis of microarrays per time-point. Heat maps are coloured by the Z-score normalised per row. Each column represents a patient. T1 is prior to the detection of HCMV DNAemia; T2 at initial detection of HCMV DNAemia; T3 is at the peak of HCMV DNAemia; T4, near the resolution of HCMV reactivation. LR, low-level HCMV reactivation; HR, high-level HCMV reactivation; SD, standard deviation.

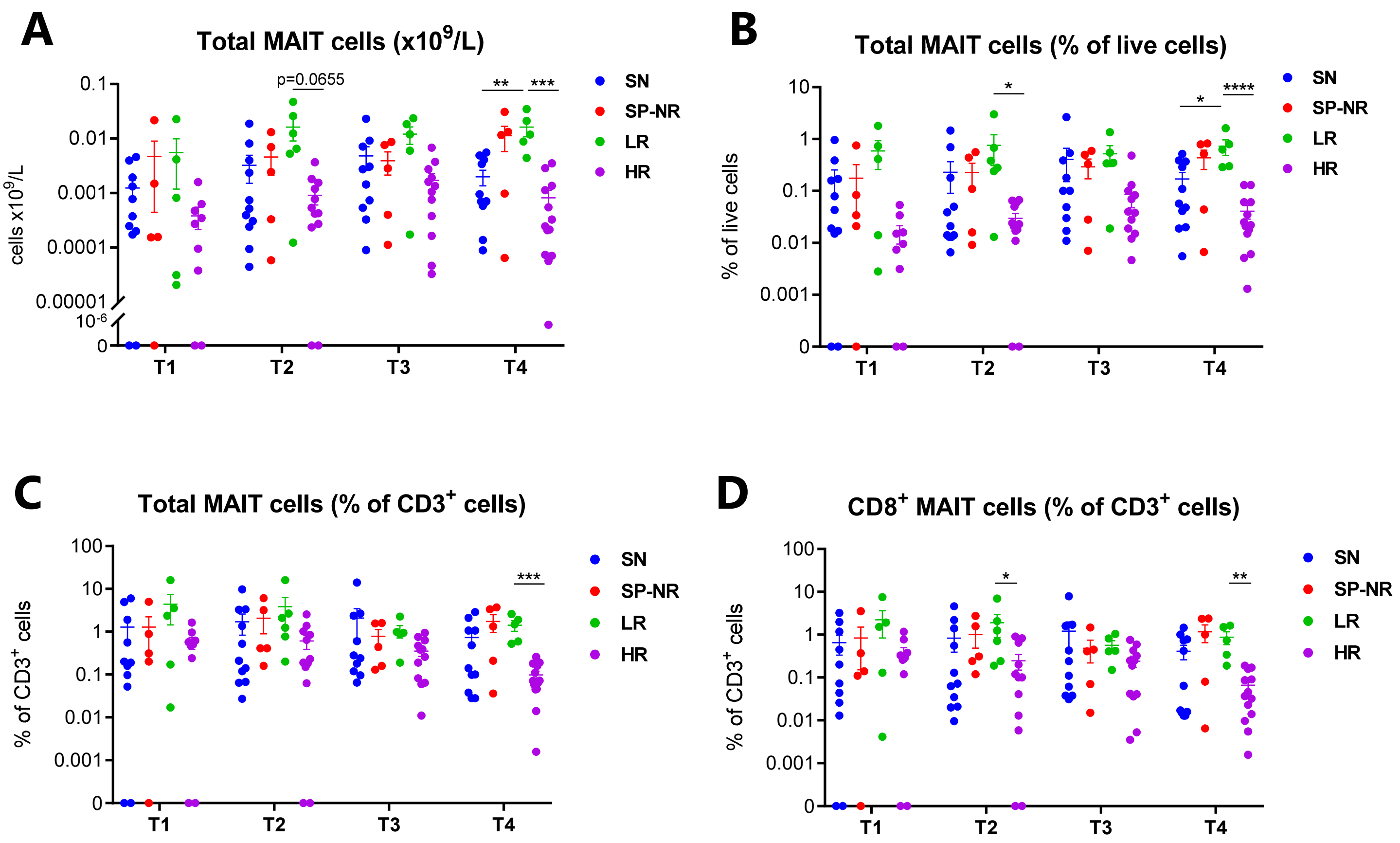


**<u>Supplemental Figure 6.</u> Total MAIT cell ($CD3^+V\alpha7.2^+CD161^{hi}$) frequencies and MAIT cells as a percentage of total $CD3^+$ PBMCs.** (**A**) Absolute counts ($x10^9$/L) of total MAIT cells ($CD3^+V\alpha7.2^+CD161^{hi}$). (**B**) Total MAIT cells ($CD3^+V\alpha7.2^+CD161^{hi}$) expressed as a percentage of total live PBMCs. (**C**) Total MAIT cells ($CD3^+V\alpha7.2^+CD161^{hi}$) expressed as a percentage of total $CD3^+$ cells. (**D**) $CD8^+$ MAIT cells ($V\alpha7.2^+CD161^{hi}$ $CD8^+$ T cells) expressed as a percentage of total $CD3^+$ cells. Mean ± SEM are indicated. Statistical significance was assessed via two-way mixed-effects models with Tukey's multiple comparisons tests (* $p<0.05$, **$p<0.01$, ***$p<0.001$, ****$p<0.0001$). SN, seronegative (blue; n=11); SP-NR, seropositive no reactivation (red; n=5); LR, low-level HCMV reactivation (green; n=6); HR, high-level HCMV reactivation (purple; n=13). T1 is prior to the detection of HCMV reactivation; T2, at the initial detection of HCMV reactivation; T3, the peak; T4, near the resolution of HCMV reactivation.

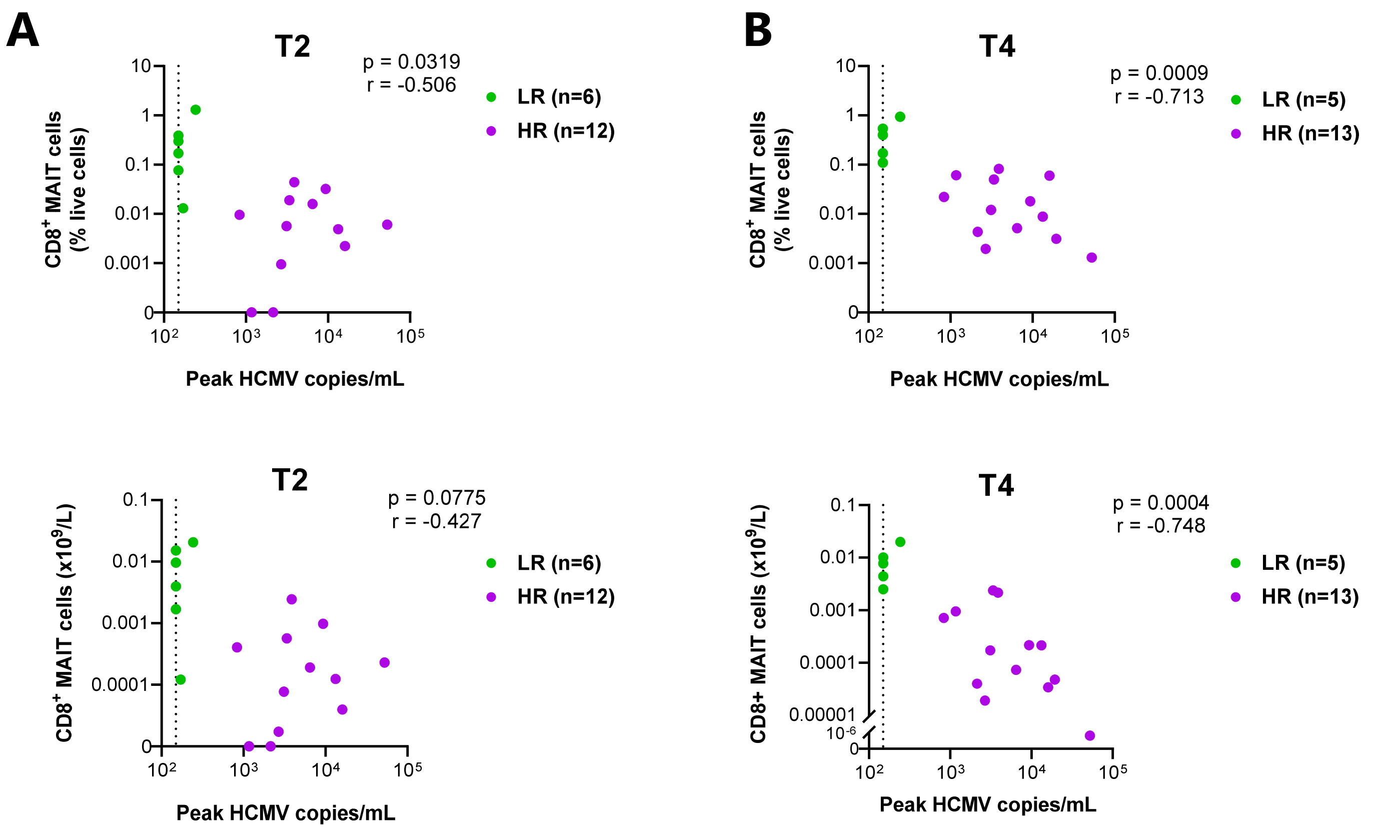


**Supplemental Figure 7. Inverse correlation between peak HCMV copies/mL and CD8$^+$ MAIT cell frequencies at T2 and T4.** Two-tailed Spearman correlations between peak HCMV copies/mL and MAIT cell frequencies at (**A**) T2 (initial detection of HCMV DNAemia) and (**B**) T4 (near resolution of HCMV DNAemia) in HSCT recipients who experienced low-level HCMV reactivation (LR; green) and high-level HCMV reactivation (HR; purple). Correlations for CD8$^+$ MAIT cell percentages (of live cells) and absolute counts (x10$^9$/L) are shown. $p<0.05$ was considered significant. The vertical black dotted line at 150 HCMV copies/mL plasma indicates the lower limit of quantitation (LLQ) of the HCMV DNA quantitative PCR assay.

**Supplemental Table 1.** **HCMV DNAemia characteristics and antiviral treatment in patients with HCMV reactivation**

| | All HCMV reactivators (n=19) | LR (n=6) | HR (n=13) | p value |
|---|---|---|---|---|
| **Day post-HSCT of initial HCMV DNAemia detection** | 29 (12-46) | 35 (12-45) | 28 (17-46) | 0.6257 |
| **Magnitude of first detected HCMV DNAemia (copies/mL)** | 150 (150-515) | 150 (150-150) | 150 (150-515) | 0.5170 |
| **Peak HCMV DNA copies/mL** | 2960 (150-52740) | 150 (150-244) | 3873 (836-52740) | **<0.0001** |
| **Duration of HCMV DNAemia (days)** | 38.0 (7-121) | 21.5 (7-49) | 60.0 (28-121) | **0.0180** |
| **HCMV DNA $log_{10}$ AUC** | | | | |
| T2 | 3.02 (2.72-3.57) | 2.72 (2.72-3.57) | 3.05 (2.97-3.50) | 0.2769 |
| T3 | 4.09 (3.20-5.35) | 3.50 (3.20-3.76) | 4.35 (3.57-5.35) | **0.0013** |
| T4 | 4.62 (3.32-5.54) | 3.68 (3.32-3.87) | 4.73 (4.29-5.54) | **0.0002** |
| Total in first 100 days post-HSCT | 4.64 (3.32-5.54) | 3.69 (3.32-3.87) | 4.74 (4.29-5.54) | **<0.0001** |
| **Antiviral therapy for HCMV reactivation (n patients)** | 14 (74%) | 1 (17%) | 13 (100%) | **0.0005** |

**Notes:** The days post-transplant to initial detection of HCMV reactivation were compared between LR and HR using a two-tailed unpaired t test. Two-tailed Mann-Whitney tests were used for other comparisons. p values indicate comparison between LR and HR groups. AUC (area under the curve), LR (low-level HCMV reactivation), HR (high-level HCMV reactivation), HSCT (haematopoietic stem cell transplant), HCMV (human cytomegalovirus). HCMV AUC not shown for T1 (as T1 is prior to the detection of HCMV DNAemia). 'Antiviral therapy' refers to pre-emptive therapy.

**<u>Supplemental Table 2.</u> Details of sample time-points for mass cytometry**

| | No HCMV reactivation | | HCMV reactivation | | |
|---|---|---|---|---|---|
| | SN (n=11) | SP-NR (n=5) | LR (n=6) | HR (n=13) | <u>All patients</u> (n=35) |
| **Number of samples (n)** | | | | | |
| T1 | 11 | 5 | 5 | 9 | 30 |
| T2 | 11 | 5 | 6 | 12 | 34 |
| T3 | 10 | 5 | 5 | 12 | 32 |
| T4 | 11 | 5 | 5 | 13 | 34 |
| **Days post HSCT** | | | | | |
| T1 | 19 (17-26) | 20 (19-25) | 19 (17-24) | 18 (14-32) | 19 (14-32) |
| T2 | 34 (33-41) | 34 (32-42) | 36 (31-45) | 33 (27-48) | 33.5 (27-48) |
| T3 | 50 (45-61) | 47 (46-61) | 47 (46-59) | 46 (39-62) | 47 (39-62) |
| T4 | 83 (75-89) | 83 (82-89) | 82 (67-87) | 84 (74-89) | 83 (67-89) |
| **Days from initial HCMV DNAemia** | | | | | <u>All reactivation patients</u> (n=19) |
| T1 | n/a | n/a | -21 (-21 to -7) | -7 (-28 to -3) | -14 (-28 to -3) |
| T2 | n/a | n/a | 0 (0-21) | 4.5 (0-19) | 3 (0-21) |
| T3 | n/a | n/a | 21 (7-42) | 21 (12-28) | 21 (7-42) |
| T4 | n/a | n/a | 56 (22-70) | 54 (37-69) | 55 (22-70) |
| **Days from peak HCMV DNAemia** | | | | | |
| T1 | n/a | n/a | -35 (-42 to -28) | -31 (-42 to -24) | -32.5 (-42 to -24) |
| T2 | n/a | n/a | -17.5 (-28 to -7) | -16.5 (-23 to -8) | -16 (-28 to -7) |
| T3 | n/a | n/a | 0 (-14 to 0) | 0 (-7 to 0) | 0 (-14 to 0) |
| T4 | n/a | n/a | 28 (13-37) | 35 (17-46) | 31.5 (13-46) |

**<u>Notes:</u>** Data are presented as absolute values (n) or median (range). SN (HCMV seronegative; D-/R-), SP-NR (HCMV seropositive no reactivation), LR (low-level HCMV reactivation; <250 peak HCMV DNA copies/mL), HR (high-level HCMV reactivation; 830-53,000 peak HCMV DNA copies/mL), HCMV (human cytomegalovirus), HSCT (haematopoietic stem cell transplant), n/a (not applicable). 'Samples' refers to PBMC samples.

**Supplemental Table 3. Mass cytometry antibody panel**

| Metal | Specificity [anti-human] | Clone | Concentration (µg/mL) | Supplier | Stain type |
|---|---|---|---|---|---|
| 89Y | CDK1 | sc-54 | 8 | Santa Cruz Biotechnology | Intracellular |
| 115In | CD11c | Bu15 | 1 | BioLegend | Surface |
| 141Pr | CD27 | M-T271 | 1 | Becton Dickinson | Surface |
| 142Nd | CD19 | HIB19 | 1 | Becton Dickinson | Surface |
| 143Nd | CD45RA | HI100 | 2 | BioLegend | Surface |
| 144Nd | CD69 | FN50 | 3 | BioLegend | Surface |
| 145Nd | CD4 | RPA-T4 | 2 | BioLegend | Surface |
| 146Nd | CD8A | RPA-T8 | 8 | BioLegend | Surface |
| 147Sm | CD20 | 2H7 | 4 | BioLegend | Surface |
| 148Nd | CD16 | 3G8 | 1 | Becton Dickinson | Surface |
| 149Sm | TIM3 | 7D3 | 1 | Becton Dickinson | Surface |
| 150Nd | TCR γ/δ | B1 | 8 | Becton Dickinson | Surface |
| 151Eu | CD278 | DX29 | 4 | Fluidigm | Surface |
| 152Sm | CD45RO | UCHL1 | 8 | BioLegend | Surface |
| 153Eu | CD304 | 12C2 | 1 | BioLegend | Surface |
| 154Sm | CD163 | GHI/61 | 6 | Becton Dickinson | Surface |
| 155Gd | CD314 (NKG2D) | 1D11 | 6 | Becton Dickinson | Surface |
| 156Gd | CD86 | IT2.2 | 2 | Becton Dickinson | Surface |
| 158Gd | CD33 | WM53 | 0.5 | Becton Dickinson | Surface |
| 159Tb | HCMV IE | 8B1.2 | 2 | Merck Millipore | Intracellular |
| 160Gd | CD14 | M5E2 | 1 | Becton Dickinson | Surface |
| 161Dy | CD274 | 29E.2A3 | 5 | BioLegend | Surface |
| 162Dy | FoxP3 | PCH101 | 4 | eBioscience | Intracellular |
| 163Dy | CD159c (NKG2C) | REA205 | 4 | Miltenyi Biotec | Surface |
| 164Dy | CD161 | DX12 | 4 | Becton Dickinson | Surface |
| 165Ho | CD127 | A019D5 | 2 | BioLegend | Surface |
| 166Er | CD34 | 581 | 4 | Becton Dickinson | Surface |
| 167Er | CD38 | HIT2 | 4 | BioLegend | Surface |
| 168Er | HCMV pp65 | B051M | 2 | Merck Millipore | Intracellular |
| 169Tm | CD25 | M-A251 | 4 | BioLegend | Surface |
| 170Er | CD3 | UCHT1 | 1 | BioLegend | Surface |
| 171Yb | Granzyme B | GB11 | 0.25 | Acris | Intracellular |
| 172Yb | CD197 (CCR7) | G043H7 | 2 | BioLegend | Surface |
| 173Yb | TCR Vα7.2 | 3C10 | 4 | BioLegend | Surface |
| 174Yb | HLA-DR | L243 | 4 | Becton Dickinson | Surface |
| 175Lu | CD279 (PD-1) | EH12.2H7 | 8 | Becton Dickinson | Surface |
| 176Yb | CD56 | NCAM16.2 | 1 | Becton Dickinson | Surface |
| 209Bi | CD57 | NK-1 | 0.5 | Becton Dickinson | Surface |

**Notes**: Antibodies were purchased either pre-conjugated from Fluidigm, or in a purified, carrier-free unlabelled format from other suppliers. Unlabelled antibodies were conjugated with the indicated metal isotope by the Ramaciotti Facility for Human Systems Biology (The University of Sydney, Australia) using the Maxpar Antibody Labelling Kit (Fluidigm). All antibodies were titrated before inclusion in the panel.

**Supplemental Table 4.** **Cell subset percentages included in SAM analysis**

| **Percentages** |
|---|
| B cells (% live cells) |
| Total monocytes (% live cells) |
| $CD16^{+}$ γδ-T cells (% γδ-T cells) |
| $CD16^{+}$ mDC (% mDC) |
| $CD16^{+}$ monocytes (% total monocytes) |
| $CD16^{+}$ NK cells (% NK cells) |
| $CD161^{low/-}$ $V\alpha 7.2^{+}$ $CD8^{+}$ T cells (% live cells) |
| $Tim3^{+}$ $CD8^{+}$ T cells (% $CD8^{+}$ T cells) |
| $CD27^{+}$ $CD4^{+}$ T cells (% $CD4^{+}$ T cells) |
| $CD27^{+}$ $CD8^{+}$ T cells (% $CD8^{+}$ T cells) |
| $CD27^{+}$ γδ-T cells (% γδ-T cells) |
| $CD27^{+}$ γδ-T cells (% live cells) |
| $CD27^{+}$ B cells (% B cells) |
| Classical monocytes (% live cells) |
| $CD3^{-}$ cells (% live cells) |
| $CD3^{+}CD33^{-}$ cells (% live cells) |
| $CD34^{+}CD3^{-}$ cells (% live cells) |
| $CD38^{+}$ $CD4^{+}$ T cells (% $CD4^{+}$ T cells) |
| $CD38^{+}$ $CD8^{+}$ MAIT cells (% $CD8^{+}$ MAIT cells) |
| $CD38^{+}$ $HLA\text{-}DR^{+}$ EM $CD4^{+}$ T cells (% $CD4^{+}$ T cells) |
| $CD38^{+}$ $HLA\text{-}DR^{+}$ EM $CD4^{+}$ T cells (% EM $CD4^{+}$ T cells) |
| $CD38^{+}$ $HLA\text{-}DR^{+}$ EM $CD8^{+}$ T cells (% $CD8^{+}$ T cells) |
| $CD38^{+}$ $HLA\text{-}DR^{+}$ EM $CD8^{+}$ T cells (% EM $CD8^{+}$ T cells) |
| $CD38^{+}$ non-classical monocytes (% non-classical monocytes) |
| $CD38^{+}GzmB^{+}$ CM $CD8^{+}$ T cells (% $CD8^{+}$ T cells) |
| $CD38^{+}GzmB^{+}$ CM $CD8^{+}$ T cells (% CM $CD8^{+}$ T cells) |
| $CD38^{+}$ $HLA\text{-}DR^{+}$ $CD4^{+}$ T cells (% $CD4^{+}$ T cells) |
| $CD3^{-}CD56^{-}CD14^{+}CD38^{+}HLA\text{-}DR^{-}$ cells (% live cells) |
| CD4:CD8 ratio |
| $CD4^{+}$ T cells (% live) |
| $CD4^{+}CD8^{+}$ T cells (% $CD3^{+}$ cells) |
| $CD45RA^{+}$ γδ-T cells (% γδ-T cells) |
| $CD45RA^{+}GzmB^{+}CD27^{-}$ γδ-T cells (% γδ-T cells) |
| $CD57^{+}$ $CD4^{+}$ T cells (% $CD4^{+}$ T cells) |
| $CD57^{+}$ $CD8^{+}$ T cells (% $CD8^{+}$ T cells) |
| $CD57^{+}$ $CD8^{+}$ T cells (% live cells) |
| $CD57^{+}GzmB^{+}$ $CD8^{+}$ T cells (% $CD8^{+}$ T cells) |
| $CD57^{+}$ NK cells (% NK cells) |
| $CD57^{+}CD38^{+}$ $CD8^{+}$ T cells (% $CD8^{+}$ T cells) |
| $CD57^{+}NKG2C^{+}$ NK cells (% NK cells) |
| $CD69^{+}$ $CD8^{+}$ MAIT cells (% $CD8^{+}$ MAIT cells) |
| $CD69^{+}$ NK cells (% NK cells) |
| $CD8^{+}$ T cells (% live) |

**Table S4** continued over page

| |
|---|
| $CD86^+$ monocytes (% total monocytes) |
| $CD86^+$ classical monocytes (% classical monocytes) |
| $CD86^+$ intermediate monocytes (% intermediate monocytes) |
| $CD86^+$ non-classical monocytes (% non-classical monocytes) |
| $CD86^+$ mDCs (% mDCs) |
| CM $CD4^+$ T cells (% $CD4^+$ T cells) |
| CM $CD4^+$ T cells (% live cells) |
| CM $CD8^+$ T cells (% $CD8^+$ T cells) |
| EM $CD4^+$ T cells (% $CD4^+$ T cells) |
| EM $CD8^+$ T cells (% $CD8^+$ T cells) |
| γδ-T cells (% live cells) |
| $GzmB^+$ $CD8^+$ T cells (% $CD8^+$ T cells) |
| $GzmB^+$ $CD4^+$ T cells (% $CD4^+$ T cells) |
| $GzmB^+$ γδ-T cells (% γδ-T cells) |
| $CD57^+GzmB^+$ $CD3^+CD56^+$ cells (% $CD3^+CD56^+$ cells) |
| HLA-$DR^+$ NK cells (% NK cells) |
| HLA-$DR^+CD38^+$ $CD8^+$ MAIT cells (% $CD8^+$ MAIT cells) |
| HLA-$DR^+CD38^+$ $CD8^+$ T cells (% $CD8^+$ T cells) |
| $ICOS^+$ $CD4^+$ T cells (% $CD4^+$ T cells) |
| $ICOS^+$ $HLADR^+CD38^+$ $CD4^+$ T cells (% $CD4^+$ T cells) |
| Intermediate monocytes (% total monocytes) |
| $CD8^+$ MAIT cells (% $Va7.2^+$ $CD8^+$ T cells) |
| $CD8^+$ MAIT cells (% live cells) |
| Total MAIT cells (% live cells) |
| $CD8^+$ MAIT cells (% $CD8^+$ T cells) |
| mDC (% live cells) |
| Naive $CD4^+$ T cells (% $CD4^+$ T cells) |
| Naive $CD8^+$ T cells (% $CD8^+$ T cells) |
| NK cells (% live cells) |
| $NKG2C^+$ $CD8^+$ T cells (% $CD8^+$ T cells) |
| $CD3^+CD56^+$ cells (% $CD3^+$ cells) |
| $CD3^+CD56^+$ cells (% live cells) |
| Non-classical monocytes (% total monocytes) |
| PD-$1^+$ $CD4^+$ T cells (% $CD4^+$ T cells) |
| PD-$1^+$ $CD8^+$ T cells (% $CD8^+$ T cells) |
| PD-$1^+$ EM $CD8^+$ T cells (% $CD8^+$ T cells) |
| PD-$1^+$ EMRA $CD8^+$ T cells (% $CD8^+$ T cells) |
| pDC (% live cells) |
| PD-$L1^+$ classical monocytes (% classical monocytes) |
| CD45RA:CD45RO ratio on $CD4^+$ T cells (% $CD4^+$ T cells) |
| EMRA $CD4^+$ T cells (% $CD4^+$ T cells) |
| EMRA $CD8^+$ T cells (% $CD8^+$ T cells) |
| $Tim3^+$ intermediate monocytes (% intermediate monocytes) |
| Tregs (% live cells) |
| $CD57^+$PD-$1^+CD38^+$ $CD4^+$ T cells (% $CD4^+$ T cells) |

### Supplemental Table 5. Absolute cell counts included in SAM analysis

| **Absolute counts ($x10^9/L$)** |
|---|
| B cells |
| $CD16^-$ NK cells |
| $CD16^+$ γδ-T cells |
| $CD16^+$ mDC |
| $CD16^+$ NK cells |
| $CD16^+$ total monocytes |
| $CD161^{low/-}$ $V\alpha7.2^+$ $CD8^+$ T cells |
| $CD27^+$ $CD4^+$ T cells |
| $CD27^+$ $CD8^+$ T cells |
| $CD27^+$ γδ-T cells |
| $CD34^+CD3^-$ cells |
| $CD38^+$ $CD8^+$ MAIT cells |
| $CD38^+$ non-classical monocytes |
| $CD38^+GzmB^+$ CM $CD8^+$ T cells |
| $CD4^+$ T cells |
| $CD45RA^+$ γδ-T cells |
| $CD45RA^+GzmB^+CD27^-$ γδ-T cells |
| $CD4^+CD8^+$ T cells |
| $CD27^+$ B cells |
| Total MAIT cells |
| $CD57^+$ $CD4^+$ T cells |
| $CD57^+$ $CD8^+$ T cells |
| $CD57^+$ NK cells |
| $CD57^+CD38^+$ $CD8^+$ T cells |
| $CD38^+$ $CD4^+$ T cells |
| $CD57^+GzmB^+$ $CD8^+$ T cells |
| $CD57^+GzmB^+$ $CD3^+CD56^+$ cells |
| $CD57^+NKG2C^+$ NK cells |
| $CD57^+PD\text{-}1^+CD38^+$ $CD4^+$ T cells |
| $CD69^+$ $CD8^+$ MAIT cells |
| $CD69^+$ NK cells |
| $CD8^+$ T cells |
| $CD8^+$ MAIT cells |
| $CD86^+$ total monocytes |
| $CD86^+$ classical monocytes |
| $CD86^+$ intermediate monocytes |
| $CD86^+$ mDC |
| $CD86^+$ non-classical monocytes |
| Classical monocytes |
| CM $CD4^+$ T cells |
| CM $CD8^+$ T cells |

**Table S5** continued over page

| |
|---|
| EM CD4$^+$ T cells |
| EM CD8$^+$ T cells |
| γδ-T cells |
| GzmB$^+$ γδ-T cells |
| GzmB$^+$ CD4$^+$ T cells |
| GzmB$^+$ CD8$^+$ T cells |
| HLA-DR$^+$ NK cells |
| HLA-DR$^+$CD38$^+$ CD4$^+$ T cells |
| HLA-DR$^+$CD38$^+$ CD8$^+$ MAIT cells |
| HLA-DR$^+$CD38$^+$ CD8$^+$ T cells |
| HLA-DR$^+$CD38$^+$ EM CD4$^+$ T cells |
| HLA-DR$^+$CD38$^+$ EM CD8$^+$ T cells |
| ICOS$^+$ CD4$^+$ T cells |
| ICOS$^+$ HLADR$^+$CD38$^+$ CD4$^+$ T cells |
| Intermediate monocytes |
| Total lymphocytes* |
| mDC |
| Total monocytes* |
| Naïve CD4$^+$ T cells |
| Naïve CD8$^+$ T cells |
| NK cells |
| NKG2C$^+$ CD8$^+$ T cells |
| CD3$^+$CD56$^+$ cells |
| Non-classical monocytes |
| PD-1$^+$ CD8$^+$ T cells |
| PD-1$^+$ CD4$^+$ T cells |
| PD-1$^+$ EM CD8$^+$ T cells |
| PD-1$^+$ EMRA CD8$^+$ T cells |
| pDC |
| PD-L1$^+$ classical monocytes |
| EMRA CD4$^+$ T cells |
| EMRA CD8$^+$ T cells |
| Tim3$^+$ CD8$^+$ T cells |
| Tim3$^+$ intermediate monocytes |
| Tregs |
| WBC* |

*derived directly from automated full blood analyser

**Supplemental Table 6. Characteristics of CD8$^{high}$ and CD8$^{low}$ HSCT recipients with HCMV reactivation**

| Characteristic | CD8$^{low}$ (n=7) | CD8$^{high}$ (n=10) | p value |
|---|---|---|---|
| **LR : HR** | 1 : 6 | 4 : 6 | 0.3382 |
| **Age (median (range))** | 47 (18 – 67) | 57.5 (18 – 62) | 0.7937 |
| **Sex (M : F)** | 2 : 5 | 4 : 6 | >0.9999 |
| ***HCMV serostatus*** | | | |
| D-/R+ | 2 (29%) | 3 (30%) | >0.9999 |
| D+/R+ | 5 (71%) | 7 (70%) | |
| ***Underlying diagnosis*** | | | |
| AML | 3 | 4 | ND |
| ALL | 1 | 4 | |
| MDS | 0 | 1 | |
| SAA | 3 | 0 | |
| MPD | 0 | 1 | |
| ***Conditioning*** | | | |
| MAC | 1 (14%) | 4 (40%) | 0.3382 |
| RIC | 6 (86%) | 6 (60%) | |
| ***Stem cell source*** | | | |
| Peripheral blood | 4 (57%) | 10 (100%) | **0.0515** |
| Bone marrow | 3 (43%) | 0 | |
| ***Donor type*** | | | |
| MUD | 4 | 7 | ND |
| Haploidentical | 0 | 1 | |
| HLA-identical related | 3 | 2 | |
| **T cell depletion** | 7 (100%) | 6 (60%) | 0.1029 |
| ***Acute GvHD*** | | | |
| Overall (grade II-IV) | 2 (29%) | 2 (20%) | >0.9999 |
| Severe (grade III-IV) | 0 | 0 | |
| **EBV reactivation** | 5 (71%) | 5 (50%) | 0.6221 |
| **Relapse** | 3 (43%) | 3 (30%) | 0.6437 |
| **Death (1$^{st}$ year post-HSCT)** | 2 (29%) | 1 (10%) | 0.5368 |
| **Death (overall follow up)** | 3 (43%) | 3 (30%) | 0.6437 |
| **Peak HCMV copies/mL** | 2690 (171-52740) | 2108 (150-16100) | 0.5999 |
| **T3 sample (days post-HSCT)** | 47 (39-62) | 46.5 (43-60) | 0.7917 |
| **T3 sample (days post initial detection of HCMV reactivation)** | 21 (7-28) | 21 (7-42) | 0.9434 |
| **HCMV $log_{10}$ AUC T3** | 4.09 (3.20-5.35) | 4.09 (3.20-4.96) | 0.7611 |
| **HCMV $log_{10}$ $AUC_{0-100}$** | 4.73 (3.69-5.54) | 4.29 (3.32-5.28) | 0.4173 |
| **Antiviral treatment for HCMV reactivation (n patients)** | 6 (86%) | 7 (70%) | 0.6029 |

**Notes:** HSCT recipients with HCMV reactivation were stratified into 'CD8$^{high}$' and 'CD8$^{low}$' groups by the presence or absence of an elevated CD8$^{+}$ T-cell dominated immune signature at the peak of reactivation (T3). Data show absolute number (n, %) or median with range. Death (overall follow up) refers to overall follow-up period of 728

(78-1242) days post-transplant (for $CD8^{high}$ and $CD8^{low}$ patients). Note that HCMV reactivators without a T3 sample (n=2) are not included. p values represent statistical comparisons between '$CD8^{high}$' and '$CD8^{low}$' groups. ALL (acute lymphoblastic leukaemia), AML (acute myeloid leukaemia), AUC (area under the curve), $AUC_{0-100}$ (HCMV DNA area under the curve in the first 100 days post-HSCT), D (donor), EBV (Epstein-Barr Virus), GvHD (graft-versus-host disease), HCMV (human cytomegalovirus), HR (high-level HCMV reactivation; 830-53,000 peak HCMV DNA copies/mL), LR (low-level HCMV reactivation; <250 peak HCMV DNA copies/mL), MAC (myeloablative conditioning), MDS (myelodysplastic syndrome), MPD (myeloproliferative disorder), MUD (matched unrelated donor), ND (not determined), R (recipient), RIC (reduced intensity conditioning), SAA (severe aplastic anaemia), T3 (peak of HCMV DNAemia).
